# Division of labor promotes the entrenchment of multicellularity

**DOI:** 10.1101/2023.03.15.532780

**Authors:** Peter L. Conlin, Heather J. Goldsby, Eric Libby, Katherine G. Skocelas, William C. Ratcliff, Charles Ofria, Benjamin Kerr

**Author notes:** Corresponding authors. (C.O.); (B.K.). These authors contributed equally to this work (Peter L. Conlin, Heather J. Goldsby).

## Abstract

Simple multicellularity evolves readily in diverse unicellular species, but nascent multicellular groups are prone to reversion to unicellularity. Successful transitions to multicellularity therefore require subsequent mutations that promote the entrenchment of the higher-level unit, stabilizing it through time. Here we explore the causes of entrenchment using digital evolution. When faced with a trade-off between cellular metabolic productivity and information fidelity, digital “multicells” often evolve reproductive division of labor. Because digital “unicells” cannot circumvent this trade-off, unicellular revertants tend to exhibit low fitness relative to their differentiated multicellular ancestors. Thus, division of labor can drive the entrenchment of multicellularity. More generally, division of labor may play a crucial role in major transitions, enriching the complexity and functionality of higher-level units while enhancing their evolutionary stability.

---

The evolution of multicellular organisms from unicellular ancestors was a profound evolutionary transition, providing the foundation for the majority of macroscopic life on this planet (*1, 2*). Multicellularity has evolved repeatedly, resulting in forms ranging from simple filaments and cellular clusters to the complex organization of differentiated tissues in animals, plants, and some groups of fungi and algae (*3*). Both the selective conditions favoring diverse forms of multicellularity and the molecular mechanisms underpinning multicellular development have received considerable attention (*4–6*). However, less studied are the reasons that multicellularity has evolved to be so stable.

Just as unicellular lineages can transition to multicellularity, multicellular lineages can revert to unicellularity given appropriate selective conditions. While many experimental studies have demonstrated that simple undifferentiated multicellularity evolves readily in diverse species (*7–10*), others indicate that these transitions can also be reversed with relative ease (*11,12*). Furthermore, experimentally evolved multicellular organisms commonly display fitness costs relative to their unicellular ancestors in the absence of selection favoring large size (*7, 8, 13*), suggesting that reversion could be selectively advantageous. If unicellular revertants can be generated and outcompete their nascent multicellular ancestors, the transition to multicellularity would be ephemeral.

Putative examples of reversion to unicellularity from multicellularity have occured in filamentous fungi (*14*), cyanobacteria (*15*), and green algae (*16*). Some multicellular lineages, however, appear more evolutionarily stable; no known free-living unicellular lineages have descended from animals, land plants, or brown algae (*3, 17*). An understanding of why multicellularity becomes entrenched rests upon determining how evolutionary reversion is avoided.

## Experimental design

We conduct computational evolution experiments to examine the conditions that promote the entrenchment of multicellularity and its mechanistic basis. We use the digital evolution software Avida (*18*), in which digital organisms are composed of either a single lower-level unit (which we refer to as a “unicell”) or multiple lower-level units (a “multicell”) (Fig. 1). Each cell includes a program (its genome), which is a set of instructions that encode all cell-level behaviors including metabolic activities and self-replication. Cells can acquire new functionality if genomic mutations appropriately modify the underlying instructions (see Tables S1-S5 for a complete list of available instructions). There are nine types of metabolic functions, each corresponding to a binary-logic task (such as calculating the bitwise **OR** of two values; Table S6). Any organism with at least one cell performing one of the nine metabolic functions can acquire resources, which are necessary for reproduction.

**Figure 1:**
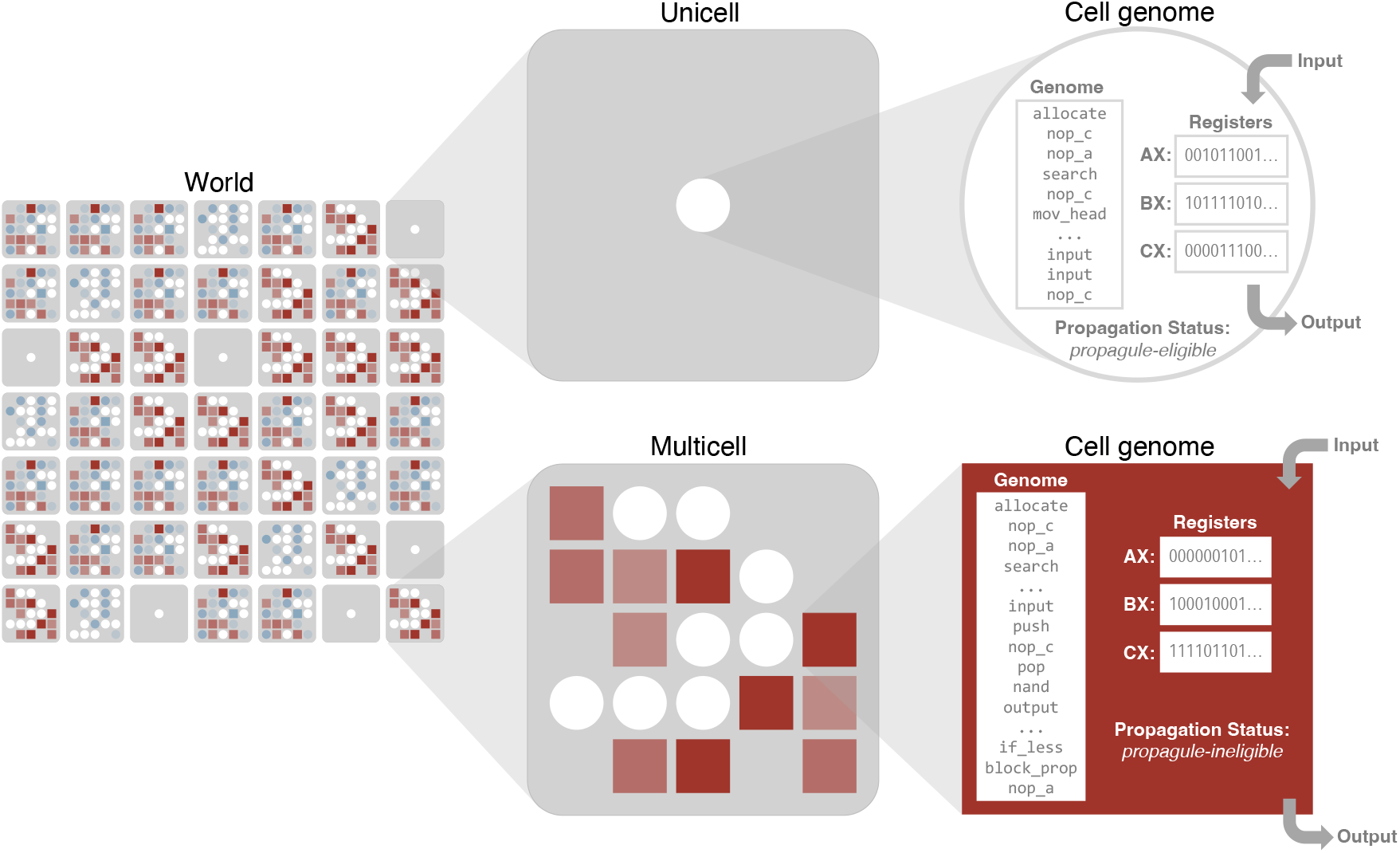
Overview of the digital evolution environment. The world comprises a set of organisms, each either unicellular or multicellular. Each cell has genomic instructions that specify the cell’s behavior, including resource gathering (via metabolic functions), intercellular communication, or cell/organism reproduction. Multicells can evolve reproductive division of labor; cells default to propagule-eligible (circles) and can initiate a new multicellular organism, but can later flag themselves “propagule-ineligible” (squares). Shading indicates the amount of dirty work performed (blue for circles, red for squares); unshaded cells (white) have not executed any mutagenic tasks.

When a cell replicates it can result in either propagule production or tissue accretion. *Propagule production* initiates a new organism. It occurs when the daughter of a reproducing cell departs from the parent organism and randomly replaces another organism within the population. *Tissue accretion* grows an existing multicell. It occurs when the daughter of a reproducing cell remains in the same body (multicell) as its parent cell. Each form of cellular reproduction is specified by distinct instructions in the cell’s genome (Table S5). Unicellular organisms must express instructions for propagule production, but not tissue accretion. A transition to multicellularity begins when mutations to a unicellular organism result in the expression of tissue accretion instructions, while retaining propagule production.

An additional distinction between unicells and multicells is the capacity for coordination. Cells within a multicell can use messaging instructions to communicate with their immediate neighbors or coordinate behaviors including terminal differentiation into a “propagule-ineligible” state. Cells in this state can divide via tissue accretion (producing propagule-ineligible daughter cells), but only propagule-eligible cells can initiate new organisms. In prior work (*19*), we found that digital multicells with these capabilities can evolve reproductive division of labor, with a metabolically-inactive germline and a metabolically-active soma, when the successful execution of a function by a cell produces mutagenic side effects in that cell’s genome. This strategy is favored because propagule-eligible (germ) cells preserve the genetic information for offspring while propagule-ineligible (somatic) cells perform the metabolic “dirty work,” increasing multicell growth rate while absorbing the mutagenic side-effects (*19–21*).

We started each evolution experiment with a digital unicellular organism (unicell) that performed one non-mutagenic function (NOT), which must be executed multiple times to produce sufficient resources for reproduction. While new functions can be introduced by mutation, the execution of mutagenic tasks can be detrimental, damaging resource acquisition or reproduction capabilities (see Table S6 for details on mutagenic levels). Each digital population evolved for one million “updates” (where cells receive, on average, 30 CPU cycles per update). Across each trajectory, we focused on the emergence of multicellularity, the evolution of mutagenic tasks, reproductive division of labor, and how these changes relate to the entrenchment of multicellularity.

## Results

### Entrenchment increases over evolutionary time

Out of 1000 evolutionary runs, 65 evolved multicellularity, including 48 that exhibited reproductive division of labor, where propagule-ineligible cells performed the vast majority of evolved mutagenic functions. These results confirm that multicellularity can evolve if collective work is beneficial and, subsequently, reproductive division of labor can be favored if the work is “dirty” (Fig. S1). For each of the 65 multicellular populations, we then measured entrenchment.

In studies of molecular evolution, a focal substitution is said to be entrenched by subsequent substitutions if it becomes relatively more deleterious to revert (*22, 23*). Here, because we are examining the entrenchment of a phenotype (multicellularity), we developed an entrenchment metric that integrates across all possible mutations that can cause reversion to unicellularity. To do this, we introduced an adjustable cost for multicellularity, implemented as additional time required for producing a propagule from a multicellular organism. During this delay, organisms remain at risk of being replaced by a reproducing competitor, but their cells are inert – they cannot perform tasks, acquire resources, replicate, or communicate. We confirmed the costly nature of this time delay by re-evolving populations from the unicellular ancestor, varying the cost of multicellularity. As expected, progressively fewer replicates evolved multicellularity at higher time delays (Fig. S2). Using this time delay cost, we measure the evolutionary stability of multicellularity by evolving populations seeded with a focal genotype under a range of increasing multicellularity costs and identify the minimum cost at which unicellular revertants dominate after 100 generations (a population is said to be dominated by unicellular revertants when the mean organism size *<* 2). “Entrenchment” is then the change in the stability of the multicellular phenotype over the period of multicellular evolution.

We measure the stability of multicellular genotypes at two time points: at the transition to multicellularity and at the end of the evolutionary run (Fig. 2A depicts a representative case study lineage). We isolated the dominant multicellular genotype at both time points and used it to initiate three test populations at exponentially increasing multicellularity costs (Fig. 2B). We see that populations of the transition genotype remain multicellular at a cost level of 64 updates, but reverted to unicellularity at or above 128 updates (Fig. 2C, yellow curve). However, for the final genotype, multicellularity persisted up to a cost level of 256 updates, but reverted at a cost of 512 updates (Fig. 2C, green curve). Thus, for this case study, multicells became entrenched during the evolution of the lineage (i.e., the change in stability from the transitional to final genotype was positive). More generally, when considering the full set of 65 multicell lineages, we saw significant entrenchment over the period of multicellular evolution (Figs. 2D, S3; Wilcoxon signed rank, *p* = 4.94*e*−12). This result raises the question: why do multicellular organisms become entrenched over evolutionary time?

**Figure 2:**
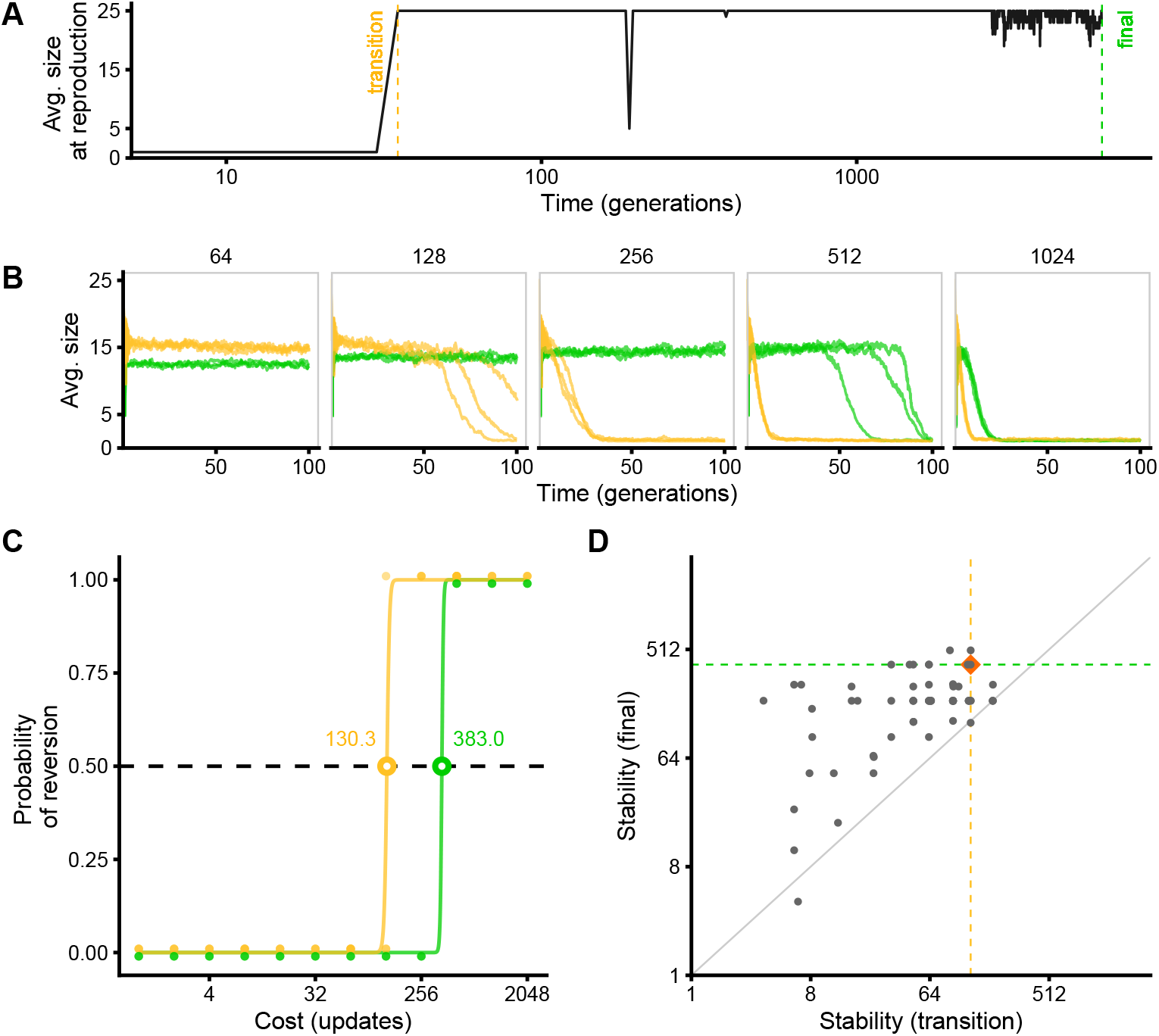
A case study for measuring entrenchment. (**A**) Size of organisms over evolutionary time (log axis to highlight early events). The time of the transition to multicellularity (yellow line) and the end of the run (green line) are indicated. (**B**) Trajectories of average organism size under different multicellularity costs, seeded with “transition” (yellow) or “final” (green) genotypes; three replicates per condition. (**C**) The probability of reversion under the full range of cost values for both the transition (yellow) and final (green) genotypes. The outcomes of each replicate are plotted as an indicator variable of the dominance of unicellular revertants. The stability value for the genotype is the cost value corresponding to 0.5 probability of reversion on a logistic fit to the data. The transition genotype has a stability value of 130.3, while the final genotype has a stability value of 383.0, indicating an entrenchment value of 252.7 (383.0 *−* 130.3). (**D**) A comparison of the stability values at the transition and final time point for all 65 multicellular lineages; for points above the diagonal, multicellularity in the lineage has become positively entrenched. The case study is indicated by the orange diamond.

### Changes in the frequency and fitness of revertants underlie entrenchment

After a transition to multicellularity, a subsequent mutation can contribute to entrenchment if it makes reversion mutations less probable or if it makes reversion mutations detrimental to relative fitness (*24, 25*). As such, we are interested in how the evolution of multicells creates a genetic background that alters the frequency and/or fitness of mutations yielding unicellular reversion. Both types of mutations can be visualized by plotting the distribution of fitness effects of reversion mutations. Entrenchment may be caused by a reduction in the number of available reversion mutations (Fig. 3A, left), a reduction in the relative fitness of revertants (Fig. 3A, middle), or both (Fig. 3A, right).

**Figure 3:**
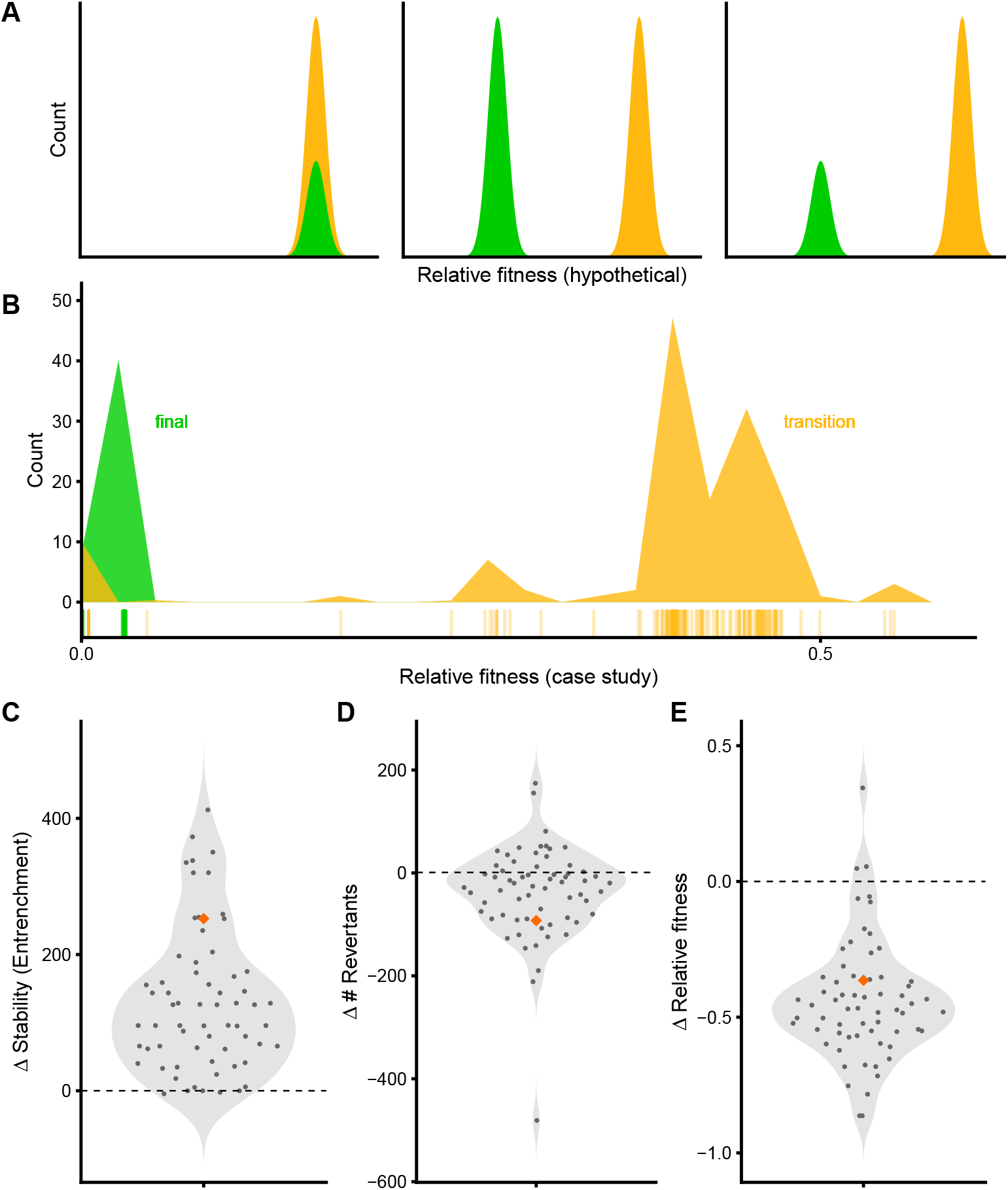
The underlying causes of entrenchment. (**A**) A schematic to illustrate three hypothetical changes in the distribution of fitness effects of reversion mutations that could cause multicellular entrenchment; yellow again indicates the distribution at the transition point, green at the evolutionary end point. Entrenchment could occur due to a decrease in the availability of reversion mutations (left), a decrease in the relative fitness of reversion mutations (middle), or both (right). (**B**) The empirical distribution of fitness effects of reversion mutations from the case study in Fig. 2 (each revertant’s fitness is also represented as a notch on the x-axis). (**C-E**) Changes in the stability value (entrenchment) and key predictor variables for all 65 genotypes that evolved multicellularity. We report the change of each value as the difference between the values at the final and transition time points. Values that correspond to the case study are indicated with an orange diamond. From left to right: (**C**) stability, (**D**) the number of unicellular revertants, and (**E**) the fitness of unicellular revertants relative to their multicellular parent.

To explore the underlying (epistatic) causes of entrenchment, we examined the evolved lineage for each multicellular replicate. We began by exploring whether evolution has altered the availability of unicellular reversion mutations. To do so, we constructed all single-locus mutants in the “transition” and “final” multicell genomes. We then tested each mutant in an environment with space for up to 32 organisms to determine if it was viable and whether it produced unicellular offspring (see Figs. S4, S5, and associated text for details). To account for stochastic effects, the mutant’s contribution to the total revertant count is the fraction of 100 replicate runs in which the mutant successfully reproduced and all descendants were unicellular. For instance, if 87 of 100 runs produced only unicellular descendants, that mutant’s contribution to its parent’s adjusted number of revertants would be 0.87. For our case study (Fig. 3B), a lineage that exhibited strong entrenchment (Fig. 3C, orange diamond), we found that there were fewer revertants for the final multicellular genotype (the integral of the green distribution) than the transitional multicellular genotype (the yellow integral). More generally, for the 65 multicellular lineages, there was a significant decrease in the number of unicellular revertants over the course of multicellular evolution (Figs. 3D, S6; Wilcoxon signed rank, *p* = 3.78*e −* 4).

We next examined whether the relative fitness effect of unicellular reversion was altered during multicellular evolution. To measure fitness, we placed a single individual in a test environment, allowed the population to grow to a maximum size of 32 individuals, and calculated the Malthusian parameter averaged over 100 replicates (excluding any replicate populations that failed to divide or re-evolved multicellularity, as described above). The relative fitness of a unicellular revertant is its Malthusian growth rate divided by that of its multicellular parent. For our case study (Fig. 3B), the revertants originating from the final multicellular genotype have lower relative fitness than revertants from the transitional genotype (the green distribution is lower than the yellow). More generally, for the 65 multicellular lineages, we see that there is a significant decrease in the relative fitness of revertants across the evolution of multicells (Figs. 3E, S6; Wilcoxon signed rank, *p* = 5.17*e −* 12). Thus, we see that the entrenchment of multicellularity (Fig. 3C) depends on two factors: evolving multicells become entrenched as unicellular revertants become both fewer (Fig. 3D) and relatively less fit (Fig. 3E).

### Germ-soma division of labor is associated with increased entrenchment

In our system, reproductive division of labor is strongly associated with entrenchment. Figure 4 explores our case study at finer resolution across its evolutionary trajectory, illustrating the relationships among multicell properties (e.g., size, workload and division of labor) and entrenchment. Reproductive division of labor evolves about midway in the trajectory and coincides with a rise in entrenchment, suggesting a potential causal link between the two. Consistent with this idea, there is a positive correlation between the change in division of labor (measured as the standard deviation of inter-cellular variance in workload across 100 replicate growth assays) and the entrenchment of evolved multicells (Fig. 5A; Spearman’s rank-order correlation, *ρ* = 0.31, *p* = 0.01). In our case study, the biggest changes in revertant relative fitness are also synchronized with the evolution of division of labor, which motivates further focus on the role that relative fitness plays in multicellular entrenchment.

**Figure 4:**
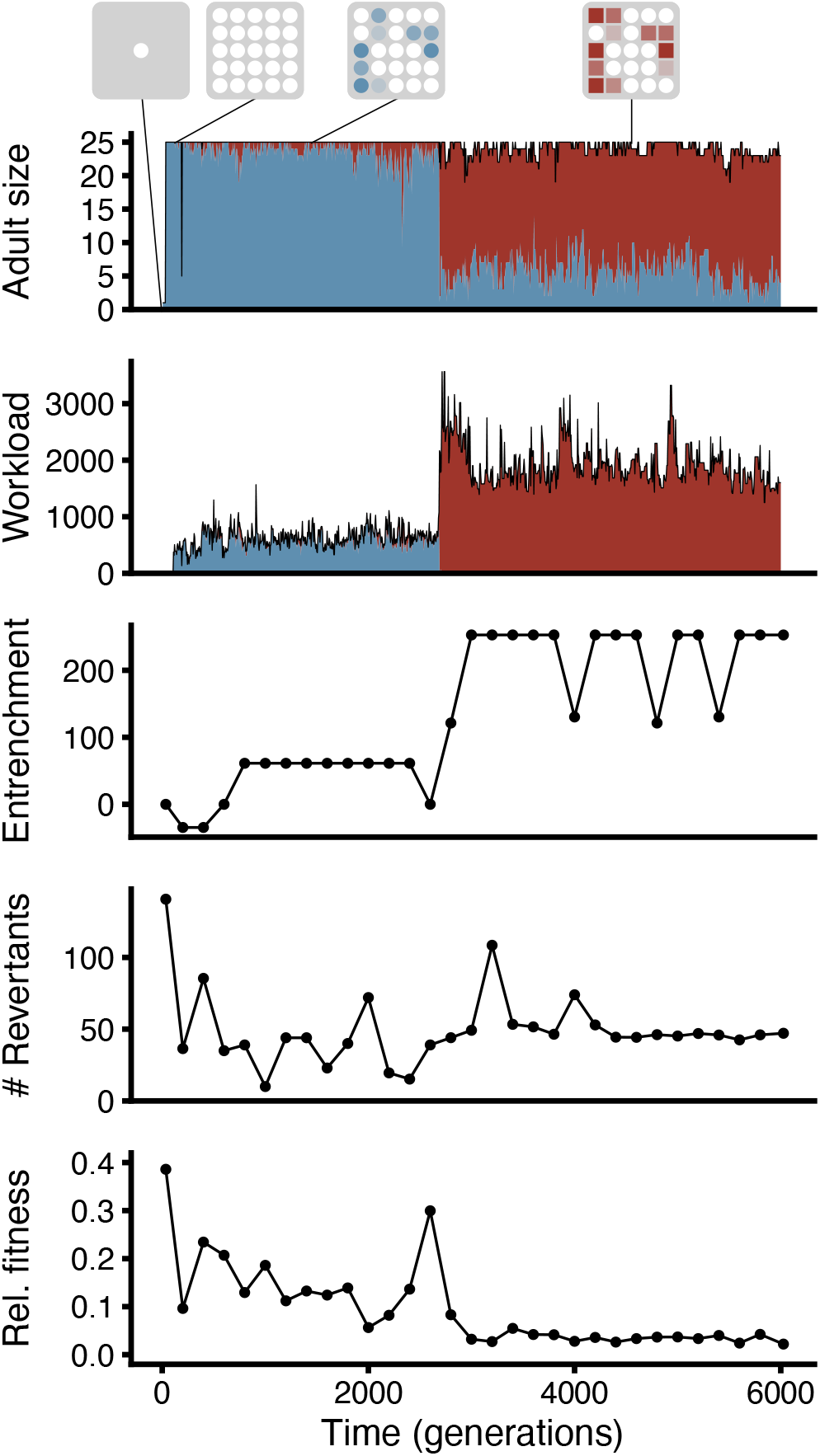
The evolutionary trajectory for our case study. Initially, organisms within this population were unicellular. Over evolutionary time, these organisms transitioned to multicellularity (as indicated by organism size, where blue represents propagule-eligible cells and red represents propagule-ineligible cells). In the early phases, most cells remained propagule-eligible and had a low workload (amount of dirty work performed). Eventually, propagule-ineligible cells evolved and began doing most dirty work while propagule-eligible cells refrained from work. Entrenchment (the change in stability of the multicellular phenotype from the transitional genotype to the genotype at the time point of interest) generally increased across the trajectory, while the number of reversion mutations decreased and the fitness of unicellular revertants relative to their multicellular parent decreased. This data supports the idea that reversion mutations become scarcer and increasingly maladaptive.

**Figure 5:**
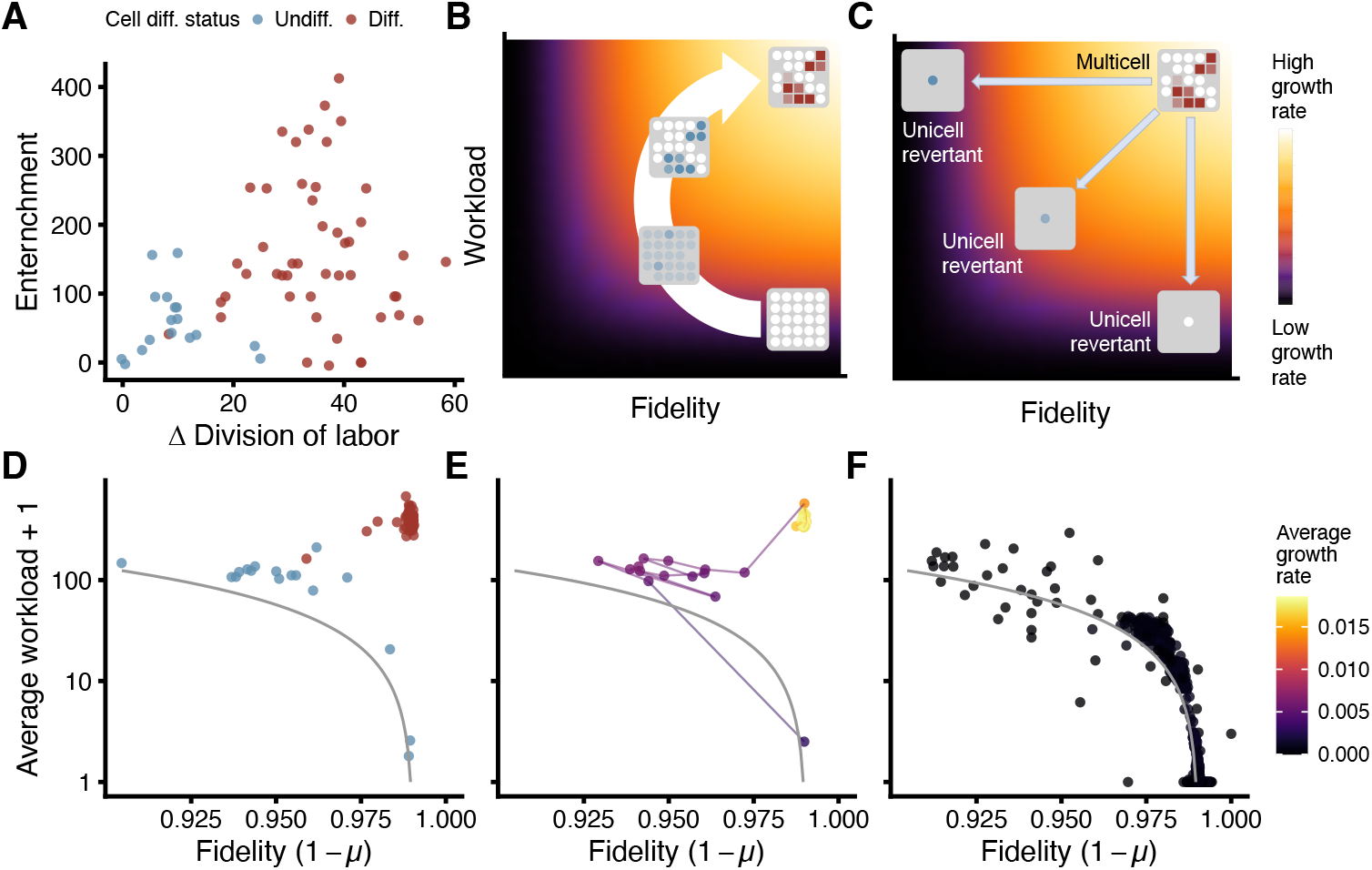
Reproductive division of labor and entrenchment. (**A**) Relationship between entrenchment and division of labor for all 65 evolved multicellular populations. Lineages that evolved reproductive division of labor (red points) were more entrenched than lineages without reproductive differentiation (blue points). (**B**) Illustration where multicellular growth rate increases at each stage along an evolutionary trajectory (large white arrow). Multicells begin with a low workload and high reproductive fidelity. Once cells evolve the capacity for mutagenic tasks, the workload rises, but the fidelity drops. Finally, reproductive division of labor allows high fidelity for germ cells and a higher workload for somatic cells. (**C**) Illustration of unicellular revertants, which face the trade-off between an active metabolism or preservation of their genetic information, and consequently realize low fitness relative to their parental multicell. (**D**-**F**) Relationship between fidelity and workload for **(D)** undifferentiated and differentiated multicell lineages at the end of the experiment, **(E)** evolution of the case-study lineage, and **(F)** all unicell revertants derived from each time point of our case study trajectory. All points are shown relative to the predicted relationship between average workload and fidelity (grey line) with fidelity = *e*^(*−N/l*)^ where *N* is a function of average workload that gives the total number of mutations and *l* is the genome length. No revertant accomplishes both high fidelity and high workload. Parts **(B), (C), (E)**, and **(F)** all use the same color scale to indicate organismal growth rate.

We find that multicellular lineages evolving reproductive differentiation (i.e., germ and somatic tissues) become significantly more entrenched than multicellular lineages without differentiation (Fig. 5A, red and blue points, respectively; Wilcoxon rank sum, *p* = 4.8*e −* 12). Furthermore, longer periods of time spent evolving as a multicell are associated with more differentiation, higher entrenchment, and lower relative fitness for unicellular revertants (Fig. S7).

We next test the idea that the evolution of reproductive differentiation allows a multicell lineage to circumvent the trade-off between metabolic work and reproductive fidelity inherent in “dirty work.” Specifically, a multicell with a metabolically active soma can achieve high resource acquisition without compromising the genetic information in its metabolically quiescent germ (Fig. 5B). However, reversion to a unicellular form reinstates the trade-off: unicell revertants can exhibit high levels of metabolic work or can preserve high fidelity genetic inheritance, but not both (Fig. 5C). Because both workload and fidelity (defined as one minus the per site mutation rate) contribute to the growth rate of a genotype, differentiated multicells have more to lose via reversion, making them disproportionately entrenched. To calculate fidelity, we directly measured the per site mutation rate (*μ*) for each evolved multicell, a time series of multicells from our case study lineage, and each unicellular revertant derived from the multicells along that same time series (see supplementary Materials and Methods for more details). As predicted by this model, differentiated multicell lineages have higher workload and higher fidelity than undifferentiated lineages (Fig. 5D). Indeed, the multicell trajectory of our case study lineage breaks free from the workload-fidelity trade-off as proposed by the model (Fig. 5E), while unicell revertants continue to be governed by this trade-off (Fig. 5F).

### Entrenchment is nullified when division of labor is disincentivized

To directly test the role of division of labor in entrenchment, we ran an experiment under conditions in which multicells could not escape the trade-off between workload and fidelity. Specifically, rather than applying the mutagenic effect of any dirty work to the cell performing the work (Fig. 6A), we applied it to the cell in that organism with the least mutagenic damage (Fig. 6B). Within this “distributed dirt” experiment, 47 out of 1000 replicates became multicellular, a proportion that is not significantly different from of our original dirty work experiment (Fisher’s exact test, *p* = 0.1). While not affecting the emergence of multicellularity, the conditions in our new experiment do remove the advantages of reproductive division of labor. Thus, we predicted that the distributed-dirt lineages would not follow the same evolutionary path as the dirty-work lineages, effectively arresting at an undifferentiated form with minimal work. Consistent with this prediction, the inter-cellular variance in workload in the distributed-dirt experiment evolved to be substantially lower than the populations from the dirty-work experiment (Fig. 6C; Wilcoxon rank sum, *p <* 2.2*e −* 16). Additionally, multicellular lineages evolved with distributed dirt did not become entrenched (Fig. S8; one-sample Wilcoxon rank sum, *p* = 0.98). Indeed, entrenchment was slightly negative in this experiment (mean distributed-dirt entrenchment value = *−*17.7), and significantly lower than both differentiated and undifferentiated lineages that evolved in the dirty-work experiment (Fig. 6D; pairwise Wilcoxon rank sum, *p <* 8.97*e −* 15).

**Figure 6:**
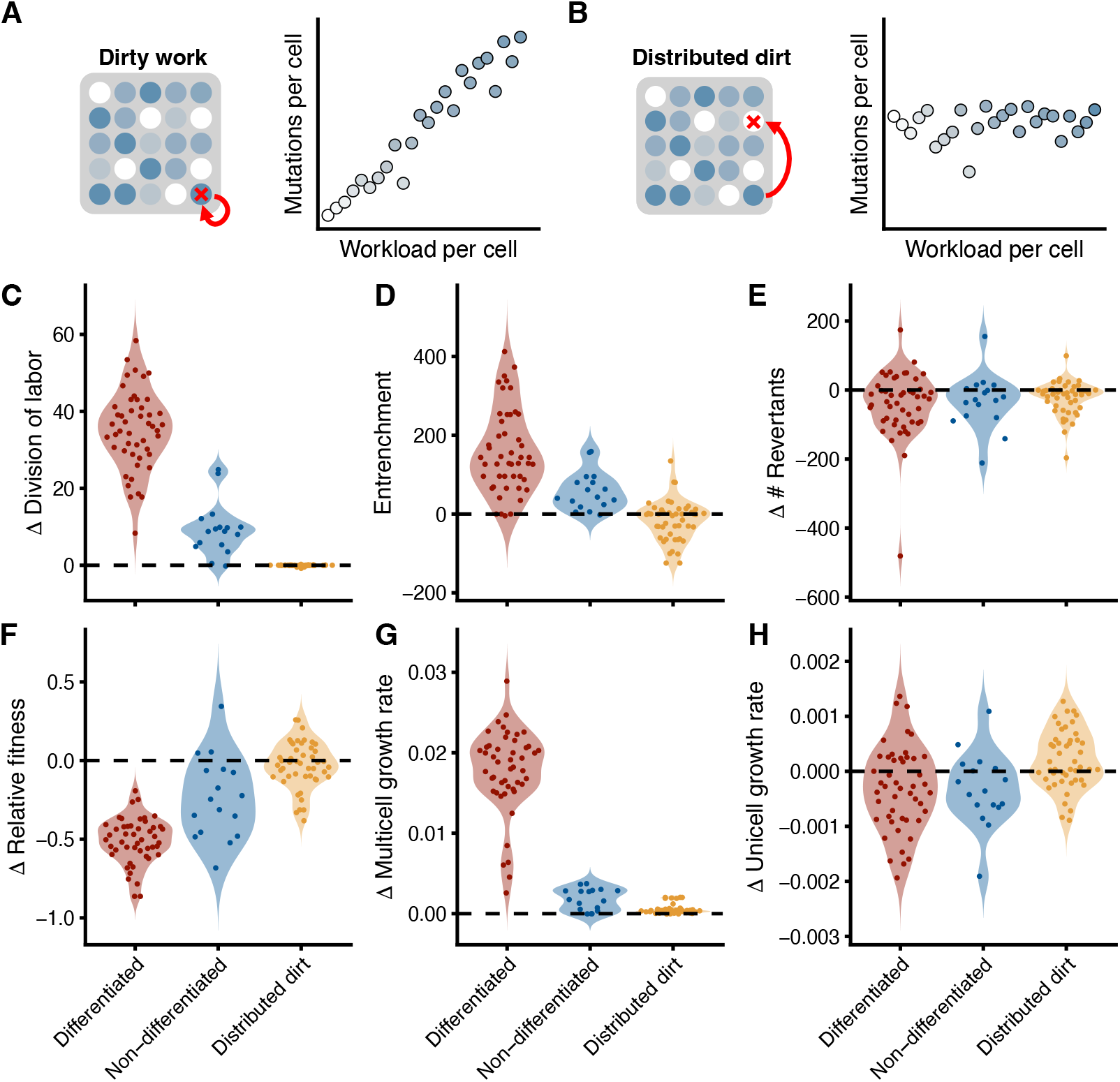
An experiment to disincentivize division of labor. (**A**) Illustration of our original “dirty work” experiment where a cell executing a task (blue circle) had mutagenic effects (red x) on its own genome. (**B**) Illustration of our “distributed dirt” experiment where the mutagenic effects are visited on the cell with the fewest prior mutations, eliminating the correlation between workload and mutations within cells. (**C**-**H**) Distributions indicating how key metrics have changed over multicell evolution for the differentiated lineages from the dirty-work experiment (red), the undifferentiated dirty-work lineages (blue), and the distributed-dirt lineages (gold). Specifically, we examine the following: (**C**) Δ division of labor (the variance in workload among cells within a multicell), (**D**) entrenchment (Δ multicell stability), (**E**) Δ number of unicellular revertant mutations, (**F**) Δ unicellular revertant relative fitness, (**G**) Δ multicell growth rate, and (**H**) Δ unicell growth rate.

Unicellular reversion mutations were generated for all distributed-dirt multicellular lineages at the transition and final time points, following the same procedures described for the dirty work experiment (see Figs. S9–S11). While we did not find significant differences in the change in the number of revertants (Fig. 6E; pairwise Wilcoxon rank sum, *p >* 0.96), there were significant differences in the change in relative fitness (Fig. 6F; pairwise Wilcoxon rank sum, *p <* 0.01) among the three types of lineages. Indeed, the change in the relative fitness of revertants was a strong predictor of the entrenchment pattern (Fig. S12; Spearman’s rank-order correlation, *ρ* = *−*0.74, *p <* 2.2*e −* 16) while changes in the number of revertants showed a weaker (albeit significant) correlation with entrenchment (Fig. S12; Spearman’s rank-order correlation, *ρ* = *−*0.21, *p* = 0.03). This is consistent with changes in revertant relative fitness being the primary driver of entrenchment.

The changes in revertant relative fitness can be further decomposed into the changes in growth rates of multicells and their unicellular revertants (Figs. 6G,H). As expected, changes in multicell growth rate negatively correlate with revertant fitness, while the opposite is true of unicellular growth rate changes (Fig. S13; Spearman’s rank-order correlation, *ρ* = *−*0.77 for multicell growth rate changes and *ρ* = 0.62 for unicell growth rate changes). Multicells exhibiting reproductive division of labor achieve the greatest gain in growth rate followed by undifferentiated multicells with moderate intercellular workload heterogeneity, while distributed-dirt multicells exhibit the smallest growth improvement (Fig. 6G; pairwise Wilcoxon rank sum, *p <* 0.01 after Bonferroni correction for all pairs). Meanwhile, the differences in the growth rate trajectories of revertants are more muted (10-fold lower change in magnitude compared with multicell). The growth rates of unicellular revertants from the differentiated and nondifferentiated multicells are negative, on average, and statistically indistiguishable from one another while the distributed-dirt unicells exhibit an average increase in growth rate (Fig. 6H; pairwise Wilcoxon rank sum, *p <* 0.01 after Bonferroni correction for pairs that include the distributed-dirt treatment).

In the differentiated dirty-work lineages, a significant decrease in revertant growth rate accompanies a significant increase in multicell growth rate. Thus, the evolution of reproductive division of labor in our system may result in a classic case of “fitness decoupling,” where the positive effects of mutations in the context of higher-level adaptation become negative at the lower level (*11, 26*). In contrast, in the distributed-dirt experiment, both multicell and revertant growth rate display significant improvement. These results suggest that in a treatment disincentivizing division of labor, mutations improving multicell fitness are more aligned with unicell fitness, leading to less entrenchment.

## Discussion

Here we have shown that the advent of cellular division of labor within nascent multicells underlies a decrease in the relative fitness of unicell revertants, thereby enhancing the evolutionary stability of multicellularity. But why does division of labor play this role? The mutations contributing to cellular differentiation involve coordination among cells occurring in the multicellular context, and thus the fitness benefits of these mutations are contingent upon prior mutations establishing multicellularity– a form of genetic epistasis (*27*). Pervasive genetic epistasis has been well-documented in protein evolution, contributing to the entrenchment of key mutations (*28–31*). However, epistasis contributing to entrenchment may be even more likely when evolutionary change causes a large reorganization of the organism, as is the case in the evolution of multicellularity (*12*). Here we see that by building upon the reorganization achieved by nascent multicells, mutations yielding division of labor epistatically constrain reversion to unicellularity.

The evolution of multicellularity is just one example of a shift in organizational complexity that occurs in any “major transition” (*1, 2*). A major transition in individuality ensues when formerly autonomous entities unite into higher-level units capable of reproduction. Drawing inspiration from two-thirds of the French tripartite motto *liberté, égalité, fraternité* - Queller introduced a revolutionary taxonomy for major transitions (*32*). The evolution of multicellularity and the shift to eusociality are examples of *fraternal* transitions, which occur when related lower-level units (e.g., insects) stay together to generate higher-level units (e.g., eusocial insect colonies). *Egalitarian* transitions result when unrelated units come together to form a higher-level unit, while lower-level units retain some reproductive rights. For example, mitochondria descend from bacteria that were engufled by an archaeal cell during an egalitarian transition into eukaryotes (*33*). Queller conceived of the “liberty” portion of the tripartite French slogan as the absence of a major transition, where lower-level units remain free-living. We think it is fruitful to extend Queller’s taxonomy by viewing “liberty” as a type of transition.

We consider a *libertarian* transition as the dissolution of a higher-level unit. Such a transition is the mirror reflection of the traditional major transition: complexity decreases and biological units jettison their nested layers. Thus, the evolutionary instability of higher-level organization in a multicellular lineage is synonymous with its vulnerability to libertarian transitions. In order to persist, higher-level entities must consistently avoid libertarian transitions through variable environmental conditions; in other words, they must become entrenched.

In line with the results in this paper, we suggest that division of labor plays a critical role in the entrenchment of different kinds of major transitions. For example, while reversion to unicellularity has occurred in relatively simple and undifferentiated multicellular taxa (*14–16*), lineages with elaborate forms of cellular differentiation have not experienced libertarian transitions to free-living unicellularity (*3, 17*). As a second example, the transition from solitary living to eusociality in various animal lineages involves reproductive division of labor at the level of individual organisms. While there have been multiple reversions to solitary living within the social bees (*34, 35*), there are no known libertarian transitions to solitary living within most eusocial taxa (including termites, ants, paper wasps, honey bees, stingless bees, and bumblebees (*34*)).

Metabolic dependencies between mutualistic partners may play a similar role in entrenching egalitarian transitions in individuality (*36*). Consider, for instance, the eukaryotic cell. Despite the fact that mitochondria and chloroplasts have retained genomes and can reproduce within the eukaryotic cell, there are no known free-living descendants of these organelles, consistent with higher-level entrenchment.

The examples above are not without exceptions. Evolutionary reversion has occurred in some eusocial lineages, such as thrips and aphids (*34*)) and transmissible cancers / cancer-derived immortal cell lineages could be viewed as a type of reversion to unicellularity (*37, 38*). Therefore, continued exploration of additional factors that contribute to higher-level entrenchment will be an important avenue for future work.

Entrenchment is an essential part of major evolutionary transitions, stabilizing emergent levels of biological organization over long time scales. The evolution of greater higher-level stability also provides insight into the nature of biological individuality: when lower-level units (e.g., cells) have evolved traits that limit their ability to survive autonomously, it is a clear signal that they have evolved into interdependent parts of a new higher-level individual (*26, 39, 40*). Here we show that entrenchment of multicellularity readily arises as a consequence of cellular specialization, inhibiting subsequent libertarian transitions in individuality. Division of labor may thus be a doubly powerful driver of increased organismal complexity, providing adaptive fuel for the evolution of large, well-integrated organisms, while simultaneously inhibiting the erosion of this complexity through evolutionary reversion.

## Acknowledgements

We thank members of the Kerr, Ofria, and Ratcliff labs for helpful feedback. Computational resources were provided by the MSU Institute for Cyber-Enabled Research.

## Authors contributions

Conceptualization: H.G., P.L.C., E.L., W.C.R., C.O., B.K.; Methodology: H.G., P.L.C., C.O., B.K.; Software: H.G., C.O.; Investigation and data curation: H.G., P.L.C.; Visualization: H.G., P.L.C., B.K.; Webbased supplementary visualizations: P.L.C., K.G.S.; Funding acquisition and administration: C.O., B.K.; Original draft: H.G., P.L.C., E.L., W.C.R., C.O., B.K.; Review & editing: All authors.

## Data and materials availability

All data and code are available on GitHub.

## Supplementary Materials

Materials and Methods

Tables S1 – S6

Figures S1 – S13

References (41 – 51)

