## Supplementary Materials for "Division of labor promotes the entrenchment of multicellularity"

#### Materials and Methods

##### Avida

Digital evolution experiments leverage the power of computers to ensure rapid generations (millions within days), perfect data collection (evolutionary lineages stored at fine temporal resolution), and high levels of experimental control (selective pressures toggled with configuration files). Here we use the software platform Avida (*1*) which has previously been employed to study the evolutionary origin of complex features (*2*), division of labor (*3, 4*), and adaptive radiation (*5*). In this study, we focus on the entrenchment of higher-level units formed via fraternal transitions, where a higher-level unit (a digital "multicell") is composed of a set of related lower-level units (digital "cells"). We maintain a population of 1,000 organisms, where each organism can be either unicellular or multicellular. Each cell consists of a program (i.e., its genome), where the instructions encode metabolism, development and reproduction.

###### Time

The standard unit of time in Avida is an "update." On average, cells receive 30 CPU cycles every update. The other unit of time in this study is a "generation." Along a line of descent, every time an offspring is produced the generation value increments by one. Within an evolving population, generation is calculated at the level of cells in the following way. Every cell gets the generation value of its parent cell incremented by one. Time in terms of generations is simply the mean generation of all cells within the population at any instant. Note that one unit of generation time within a population will track the average population turnover time of unicellular organisms, but will only be a fraction of the turnover time for multicellular organisms.

##### Instructions and functions

The Avidian genome is composed of instructions that enable cells to acquire resources, communicate with other cells in a multicellular context, reproduce within the body, and determine their eligibility to produce a propagule to initiate a new organism. A complete list of the instructions used in our evolution experiments, separated by category, are provided in Tables S1-S5.

| Instruction | Description |
| --- | --- |
| <nop_a> | No−operation instruction. Does nothing when executed, but may alter the behavior of other instructions it follows. |
| <nop_b> | No−operation instruction. Does nothing when executed, but may alter the behavior of other instructions it follows. |
| <nop_c> | No−operation instruction. Does nothing when executed, but may alter the behavior of other instructions it follows. |
| <nop_x> | No−operation instruction. Does nothing when executed and will NOT change the behavior of other instructions. |

Table S1: **No-operation instructions.** Instructions used in this study that have no direct effects when executed.

| Instruction | Description |
|---|---|
| <allocate> | Allocates extra memory for the organism to copy it's genome into. This genome can be used for either tissue accretion or propagule formation. Appears in source code as <h_alloc>. |
| <copy> | Duplicates the instruction pointed to by the read head into the position pointed to by the write head, then increments both heads. This duplication process is subject to mutations. Appears in source code as <h_copy>. |
| <search> | Identifies other positions in the genome based on complimentary patterns of nop instructions. Appears in source code as <h_search>. |
| <if_less> | Compares two values in memory. Executes the next instruction if the first value is less than the second; otherwise skips it. |
| <if_equal> | Compares two values in memory. Executes the next instruction if the first value equals the second; otherwise skips it. |
| <if_not_equal> | Compares two values in memory. Executes the next instruction if the first value does not equal the second; otherwise skips it. |
| <if_label> | Reads the series of nops that follow this instruction. If a complementary series of nops was just copied, executes the next instruction, otherwise skips it. |
| <jump_head> | Reads a value from memory and shifts the position being executed in the organism's genome by that amount. |
| <mov_head> | Moves heads (read, write, or execution) to the position of the flow head in the genome. |
| <pop> | Retrieves a number from the stack. |
| <push> | Stores a number on the current stack for later use. |
| <swap> | Swaps two values in the organism's memory. |

Table S2: **Hardware control instructions.** Instructions used in this study that control the reading/writing of the Avidian genome and handle memory.

| Instruction | Description |
|---|---|
| <dec> | Decreases a value in memory by one. |
| <inc> | Increases a value in memory by one. |
| <nand> | Performs a bitwise nand ("not−and") instruction on two values in the organism. |

Table S3: **Math instructions.** Instructions used in this study that execute mathematical operations.

| Instruction | Description |
|---|---|
| <broadcast_message> | Sends a message to all neighboring cells. Appears in source code as <bc_msg>. |
| <broadcast_message_check_task> | Packages up two numbers from the cell's registers as a message and sends them to all the neighboring cells of the focal cell. Also checks if the message contains a task result. If so, the cell performs that task. Appears in source code as <bc_msg_check_task>. |
| <donate_res_to_organism> | Gives the resources collected by the cell to the organism proper. This must be done for the organism to be able to use resources for propagule production. Appears in source code as <donate_res_to_group>. |
| <input_two_values> | Takes in two numbers as input (these inputs are fixed). Appears in source code as <fixed_input>. |
| <is_neighbor> | Sets a value in memory to one or zero based on whether the grid cell faced is occupied or not. |
| <output> | Outputs a number to be evaluated for task completion. |
| <rotate> | Rotates the facing of a cell clockwise by a number of steps equal to a value in memory. Each step is a 45°. Negative values rotate counter−clockwise. |
| <rotate_right_one> | Rotates the facing of a cell clockwise by 45°. Appears in source code as <rotate_cw>. |
| <rotate_left_one> | Rotates the facing of a cell counterclockwise by 45° (cells are embedded within a regular square lattice with neighbors in the cardinal directions). Appears in source code as <rotate_ccw>. |
| <retrieve_message> | Reads a message (if any has been received) by putting the two numbers in the message into the cell's registers. Appears in source code as <rx_msg>. |
| <send_message> | Packages up two numbers from the cell's registers as a message and sends them to the neighboring cell that the focal cell is facing. Appears in source code as <tx_msg>. |
| <send_message_check_task> | Packages up two numbers from the cell's registers as a message and sends them to the neighboring cell that the focal cell is facing. Also checks if the message contains a task result. If so, the cell performs that task. Appears in source code as <tx_msg_check_task>. |

Table S4: **Interaction instructions.** Instructions used in this study that enable communication and interaction among cells within a multicellular organism.

| Instruction | Description |
|---|---|
| <block_propagation> | Marks the cell as propagule ineligible. All cells start propagule eligible. The status is inherited from a parent cell to its daughter. Once executed, there is no way for a cell (or its descendants) to reverse this status in order to become propagule eligible again. Appears in source code as <become_soma>. |
| <produce_tissue> | Produces a new cell in the same organism. Works using the standard Avida copy loop. The daughter cell is placed in a neighboring space that the parent cell is facing if it is empty (i.e., cells cannot overwrite other cells). Appears in source code as <h_divide_local>. |
| <produce_propagule> | Produces a propagule if sufficient resources have been collected and the copy loop has executed properly. A propagule eligible cell is selected at random from the organism. It is copied to initiate a new organism with this single propagule cell. The new organism replaces an organism selected at random within the population. Appears in source code as <h_divide_remote>. |
| <if_propagule_eligible> | Executes the next instruction if the cell is propagule eligible. Appears in source code as <if_germ>. |
| <if_propagule_ineligible> | Executes the next instruction if the cell is propagule ineligible. Appears in source code as <if_soma>. |
| <if_res_less_than_threshold> | Executes the next instruction if the organism has fewer resources than are required for propagule production. Appears in source code as <if_res_less_than_thresh>. |
| <if_res_more_than_threshold> | Executes the next instruction if the organism has more resources than are required for propagule production. Appears in source code as <if_res_more_than_thresh>. |

Table S5: **Biological instructions.** Instructions used in this study that underpin cellular differentiation, multicellular development, and reproduction.

The metabolism of digital cells involves the completion of logic functions, where the execution of a series of genomic instructions enables the cell to export the result of a logic operation on input bitstrings. The nine possible logic functions that can be performed are NOT, NAND, AND, ORNOT, OR, ANDNOT, NOR, XOR, and EQUALS. The execution of each function enables the cell to sequester resources from a virtual resource pool, where there is one resource pool per organism per function. Each resource pool has a constant in-flow rate of one unit per update (the standard unit of time in Avida), while at the same time 1% of the available resources flow out, limiting total accumulation to 100 units. When a cell exports the result of a function, it uptakes 5% of the available resource associated with that function. For each function, we also specify a per-site probability of mutation each time a function is performed, referred to a "function mutagen level" (FML). Functions NOT and NAND are always non-mutagenic and thus have an FML of 0. For the treatments described in this work, the FML of the remaining seven functions increases with the complexity of the function (see Table S6). The resources acquired by successful execution of functions are required for an organism to reproduce.

| Function | FML |
|---|---|
| NOT | 0.0 |
| NAND | 0.0 |
| AND | 0.00075 |
| ORNOT | 0.00150 |
| OR | 0.00225 |
| ANDNOT | 0.00300 |
| NOR | 0.00375 |
| XOR | 0.00450 |
| EQUALS | 0.00525 |

Table S6: **The nine logic functions are listed in order of their complexity.** The probability that a genomic instruction is mutated upon execution of a function (i.e., the function mutagen level, or FML) increases incrementally with functional complexity.

**Workload**

A cell's workload is defined as:

$$w = \sum_{i \in F} N_i \frac{FML_i}{FML_{AND}}$$

where $F \equiv \{$NOT, NAND, AND, ORNOT, OR, ANDNOT, NOR, XOR, EQUALS$\}$, $N_i$ is the number of times function $i$ is performed, $FML_i$ is the mutagenic level of function $i$ (as specified in Table S6), and $FML_{AND}$ is the base $FML$ (i.e., the mutagenic level of the function AND). We define propagule workload difference as the mean workload of propagule-ineligible cells minus the mean workload of propagule-eligible cells.

**Mutations**

Mutations to the genome, in which one instruction is replaced by another randomly selected instruction, occur during two different processes: organismal reproduction and the execution of "dirty work" (logic tasks with FML>0). Mutations during reproduction only occur during propagule production (not tissue accretion) with probability 0.01 per site. On average this results in one new mutation per propagule produced. Dirty work mutations occur immediately after a cell performs a task associated with a strictly positive function mutagen level. Each site in the genome of the performing cell experiences mutations with probability given by the FML of the task executed (Table S6). More details behind the dirty work framework are provided in Goldsby et al. 2014 (*4*).

**Baseline evolution experiment**

The ancestral organism is unicellular with a genome of 100 instructions. Execution of the ancestral genome initially enables only a few simple behaviors. Specifically, the ancestor completes the task NOT a single time, subsequently rotates counterclockwise and then tries to produce a propagule cell. This reproductive process fails since it only has collected 5 units of resource. It repeats this execution loop until it has enough resources to produce a propagule (taking roughly 1800 updates for the first generation). The population has a maximum size of 1000 organisms and evolves for a total of 1 million updates with a reproductive mutation probability of 0.01 and a dirty work mutation probability defined by whatever mutagenic functions evolve in the organisms.

#### Evolution of multicellularity

Within our system, multicellularity is generally beneficial to organisms as a result of the increased ability of multiple cells to rapidly access resources that can be devoted to propagule production. Here we explore how the efficiency of resource consumption affects the evolution of multicellularity. Within our original experiments, each cell was able to consume 5% of the available resources each time a task was performed regardless of the number of cells in the organism. Here, we observe what happens when we increase the efficiency of resource uptake from 5% to 8%. This efficiency is applied at the cell level and thus is available to cells within either a unicellular or multicellular context. As can be seen in Figure S1, increasing efficiency increases the percentage of replicates that evolve to be multicellular.

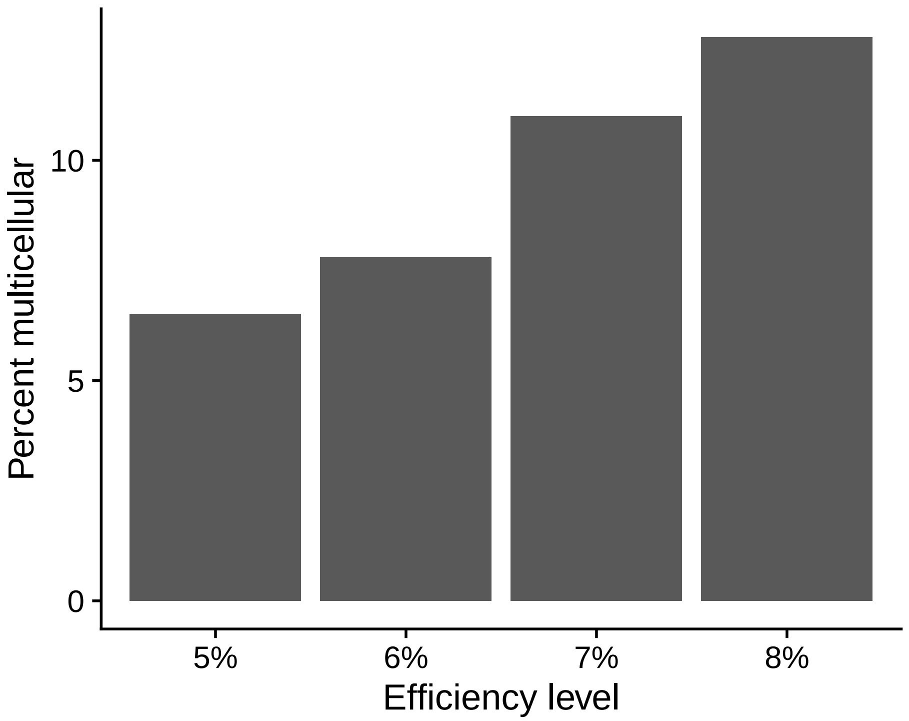


Figure S1: **Percent of evolutionary runs that evolve multicellularity as a function of resource acquisition efficiency.** For each treatment, depicted along the x-axis, we change the efficiency with which cells are able to acquire resources, and thus the benefits of multicellularity. Our original treatment (5%) has 1,000 replicates. The other treatments have 500 replicates. As the efficiency of resource consumption increases, so does the percentage of runs that evolve to be multicellular.

#### Cost of multicellularity

We added an extrinsic cost of multicellularity for the purposes of testing how readily populations would revert to back to unicells. This cost was implemented as a time delay for producing a propagule from a multicellular organism. During this delay the organism and all its constituent cells are inert – they cannot perform tasks, acquire resources, reproduce, or communicate. Additionally, the organism remains at risk for being replaced by a reproducing competitor.

To confirm the costly nature of this time delay, we reran the evolution of our initially unicellular digital populations varying this delay from 0 updates (as in our original treatment) to 512 updates. We varied this multicellularity cost from 0 updates (as in our original evolution treatment) to 512 updates, incrementing by powers of 2 (e.g., 0, 1, 2, 4, 8, 16, etc.). As expected, fewer replicates evolved multicells as this cost was increased (Fig. S2).

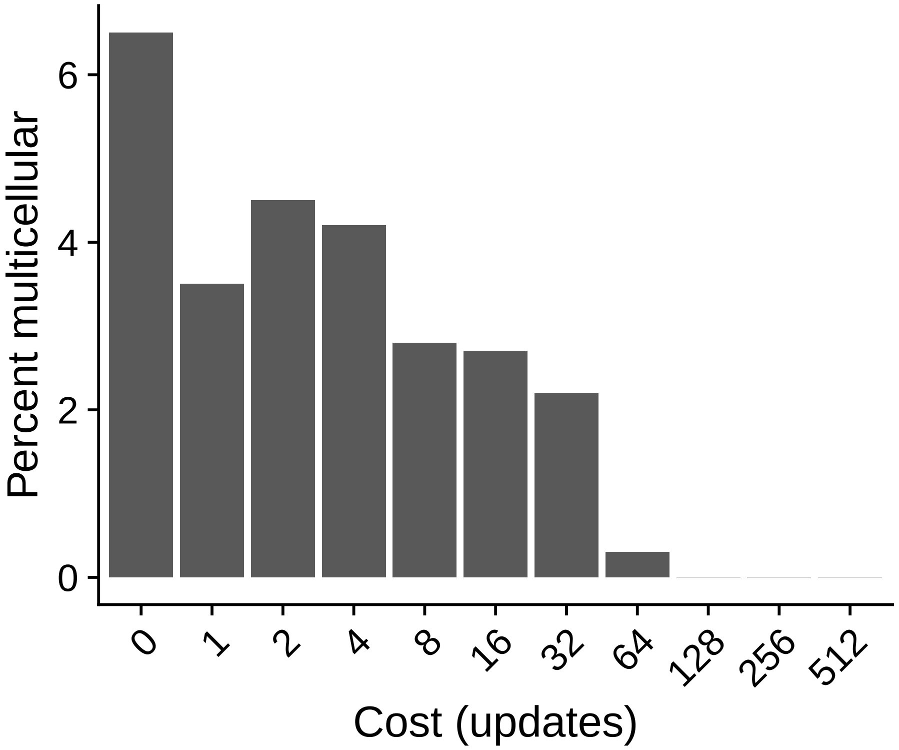


Figure S2: **Percent of evolutionary runs that evolve multicellularity as a function of time delay cost.** For each treatment, depicted along the x-axis, we change the cost of propagule production, and thus the benefits of multicellularity. We ran 1000 replicates of each treatment. The percentage of runs that evolve multicellularity decreases as the time delay cost increases.

#### Entrenchment

In studies of molecular evolution, a focal substitution is said to be entrenched by subsequent substitutions if it becomes relatively more deleterious to revert as a result of those subsequent substitutions (*6–9*). Here, because we are examining the entrenchment of a *phenotype* (multicellularity), we require an entrenchment metric that integrates across all possible mutations that can cause reversion to unicellularity. Specifically, we assess the degree to which multicellularity has become entrenched, by performing the following steps. First, we create a new population with 1000 copies of the organism when it was born. We then add an externally imposed cost of multicellularity. We run the population with this new multicellularity cost and observe if it reverts to unicellularity within 100 generations (assessed by measuring whether mean organism size has dropped below 2 cells after 100 generations of growth). We repeat this measurement with the following multicellularity cost values: 1, 2, 4, 8, 16, 32, 64, 128, 256, 512, 1024, 2048. To account for the stochastic nature of this assay, especially given the effect of dirty work on the organisms, we repeat the measurement three times for each organism, cost, and time point. We then compute a stability value for each genotype as the multicellular cost at which the probability of reversion to unicellularity $= 0.5$ using a logistic fit to the data (Fig. S3). Finally, entrenchment is computed as the difference in stability value at the final and transition time points.

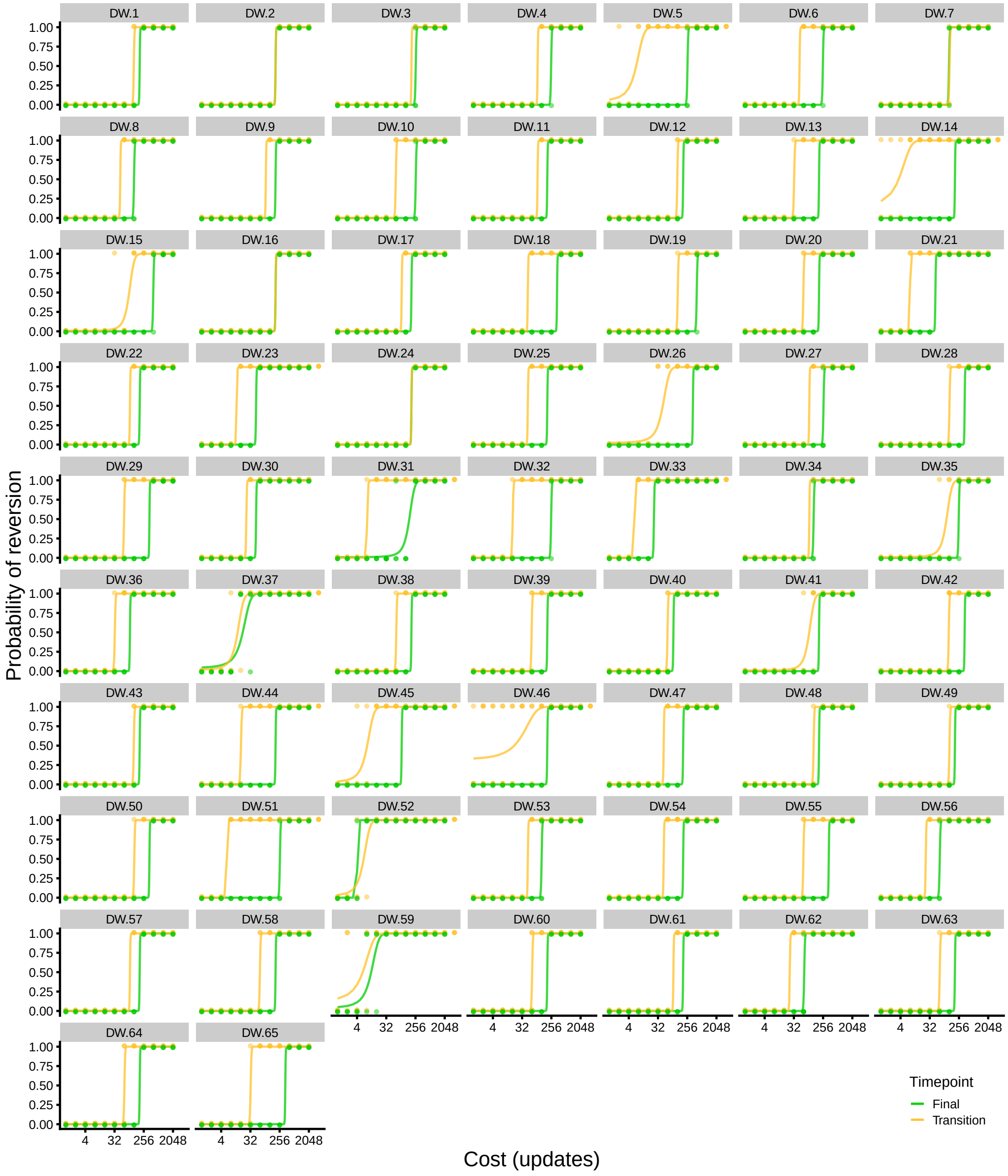


Figure S3: **A comparison of the stability of multicellularity at the transition and final time points of our "Dirty Work" evolution experiment.** The stability of multicellularity was measured for each of the 65 genotypes that evolved multicellularity at the transition (yellow) and final (green) time points of our experiment. The full range of cost values is shown with the outcomes of each replicate plotted as an indicator variable of the dominance of unicellular revertants. Logistic fits of the indicator variable are shown in yellow and green, representing the transition and final time points, respectively.

#### Generation and assessment of unicellular revertants

To understand the forces that govern whether or not a multicell is entrenched, we create all possible single-locus mutants at two time points: the "transitional" multicellular organism along the line of descent of the dominant multicell and "final" multicellular organism within this same lineage. For each single-locus mutant, we assess its viability as well as its fitness.

The first step in the assessment pipeline is the viability assay, which is a simple test to rule out organisms that are nonviable from further computationally expensive assessment. To assess a mutant's viability, we run the mutated genome for an extended period of time (up to 10,000 updates) without mutations occurring as the result of dirty work. If, within this time period, the mutated organism is able to accumulate the resources required for reproduction and to execute the instruction to create a new organism, we consider it viable and stop the viability assay. Viable mutants are then further classified as either multicellular (indicating that it executed tissue accretion instructions resulting in cellular reproduction and thus contains more than two cells) or unicellular.

This viability assay is used as a gateway to further screening and thus its conclusions are refined by the subsequent growth assay. For example, since the organism is running its code in isolation it does not actually produce an offspring organism. Additionally, it may not contain propagule-eligible cells and thus may not be able to produce offspring when grown in a population under standard conditions. If so, later assays will classify it as nonviable. Similarly, the organism may be unicellular at the time of its first successful execution of the propagule production instruction and may later execute tissue accretion instructions, becoming multicellular. If so, later assays will classify it as multicellular.

##### Growth assay and measuring relative fitness

To measure the fitness of a multicellular progenitor strain or a unicellular revertant strain, we create a solitary organism with the genome of interest and place it into an empty population with room for 62 organisms. The organism is allowed up to 50,000 updates to reach a population size threshold of 32 organisms. Once this threshold is reached, or exceeded (multiple simultaneous reproduction events can push populations above the 32 organism threshold), we calculate the average rate of increase, or Malthusian parameter, $M$, as

$$M = log(N_f/N_i)/T \tag{1}$$

where $N_f$ and $N_i$ are the final and initial number of organisms in the population and $T$ is the time it took the organism to fill the entire population. In the case that the organism fails to fill the entire population by the end of the growth assay, $T$ is set to the maximum allowed time of 50,000 updates.

To account for the effects of mutation, we repeat this process 100 times for each unicell revertant genotype and its multicell progenitor strain with different random number "seeds," leading to a different set of mutations and different cell behaviors. The fitness of a given unicell revertant relative to its multicell progenitor is defined as the ratio of their average Malthusian parameters across these replicate measurements.

##### Identification of unicellular revertants

Some mutations that occur over the course of the growth assay could result in data that may no longer be suitable for our analyses. For example, mutations that cause an organism to re-evolve multicellularity can occur during the growth assay, confounding our ability to accurately measure the relative fitness of *unicellular* revertants. Another potentially confounding scenario is that the organism we initiate the population with may fail to divide in the allotted time. This can occur if the initial organism flags itself as "propagule ineligible" or if it incurs a lethal mutation, as a result of dirty work, prior to the first propagule production event. To avoid these potentially confounding effects, we employ a two-step data filtering pipeline.

We begin by assessing whether the organisms in the population have remained unicellular for the entire duration of the growth assay. To do this, we check that the total number of cells in the population at the end of the growth assay is not greater than the total number of organisms. In total, 42.5% of our growth rate assays are "contaminated" with re-evolved multicells. We see considerable diversity within our unicell revertant genotypes; a single unicell genotype can produce a variety of outcomes: some which remain unicellular and others which re-evolve multicellularity. This is fully in line with our expectation that different mutations will occur in organisms started with different random seeds. While some multicellular progenitor genotypes appear to produce revertants that re-evolve multicellularity at higher rates than others, we find that the probability distributions for re-evolving multicellularity are quite similar at the transition and final time points (Fig. S4).

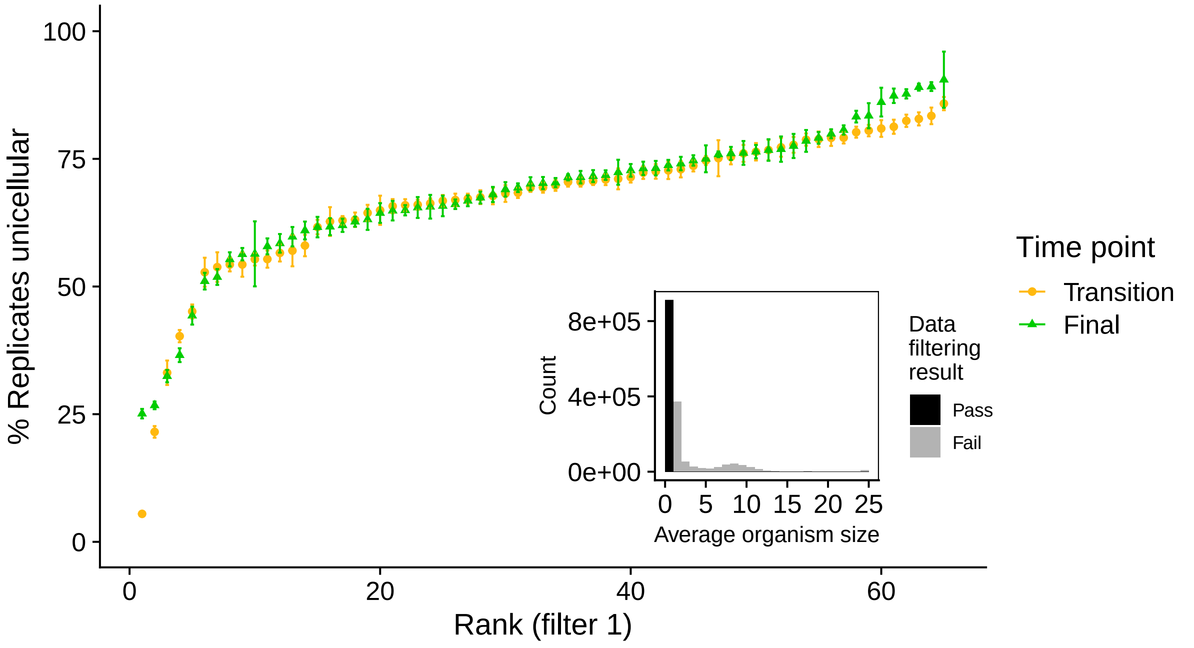


Figure S4: **Filtering unicellular revertants from the "Dirty Work" experiment that re-evolve multicellularity.** The mean and standard error for the percent of replicate populations that remain unicellular for the transition (yellow) and final (green) time points of our experiment. Unicellular revertant genotypes are grouped according to their multicellular parental genotypes and ranked from least to most replicates that remain unicellluar for each time point independently. (**Inset**) Schematic representation of our filtering criterion using data aggregated across all genotypes and both time points. We filter out any population in which the average organism size exceeds 1, indicating that multicellularity has re-evolved. Grey color indicates replicates that fail to pass the filter, black indicates those that do.

We next determine whether the organisms used to initiate the population have successfully reproduced at least once in the allotted 50,000 generations. To do this, we simply check that the total number of cells in the population is greater than 1. In total, 19% of our remaining replicate populations fail to successfully reproduce. As above, we find that some multicell progenitor genotypes are more likely than others to produce unicellular revertants that fail to successfully reproduce. However, in this case, we find that multicells from the final time point are more likely to produce inviable unicellular revertants than transition multicells (Fig. S5).

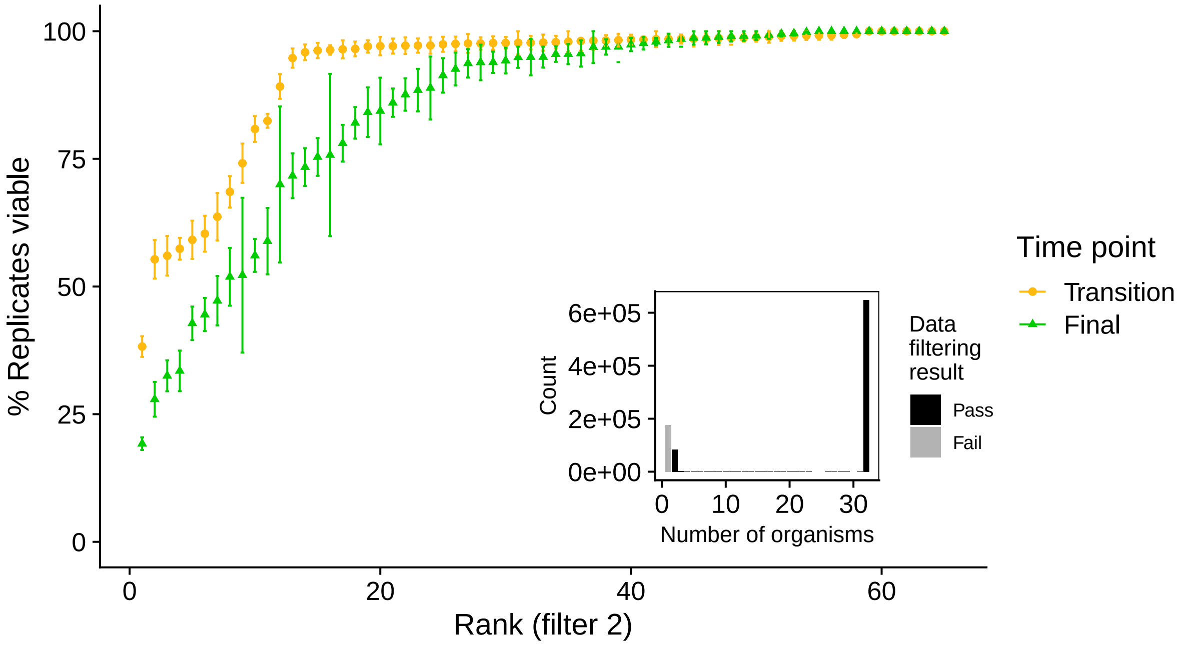


Figure S5: **Filtering unicellular revertants from the "Dirty Work" experiment that fail to reproduce.** The mean and standard error for the percent of replicate populations that successfully reproduce for the transition (yellow) and final (green) time points of our experiment. Unicellular revertant genotypes are grouped according to their multicellular parental genotypes and ranked from least to most replicates that remain unicellluar for each time point independently. (**Inset**) Schematic representation of our filtering criterion using data aggregated across all genotypes and both time points. We filter out any population in which the number of organisms at the end of the growth assay is less than 2, indicating that the organism that seeded the population has failed to reproduce. Grey color indicates replicates that fail to pass the filter, black indicates those that do.

#### Distribution of fitness effects of unicellular reversion mutations

Using only the runs that passed both of our quality control filters (see text above), we then explored the distribution of fitness effects of reversion mutations for each multicellular lineage at the time of the transition to multicellularity and at the final time point of the evolutionary run (Fig. S6).

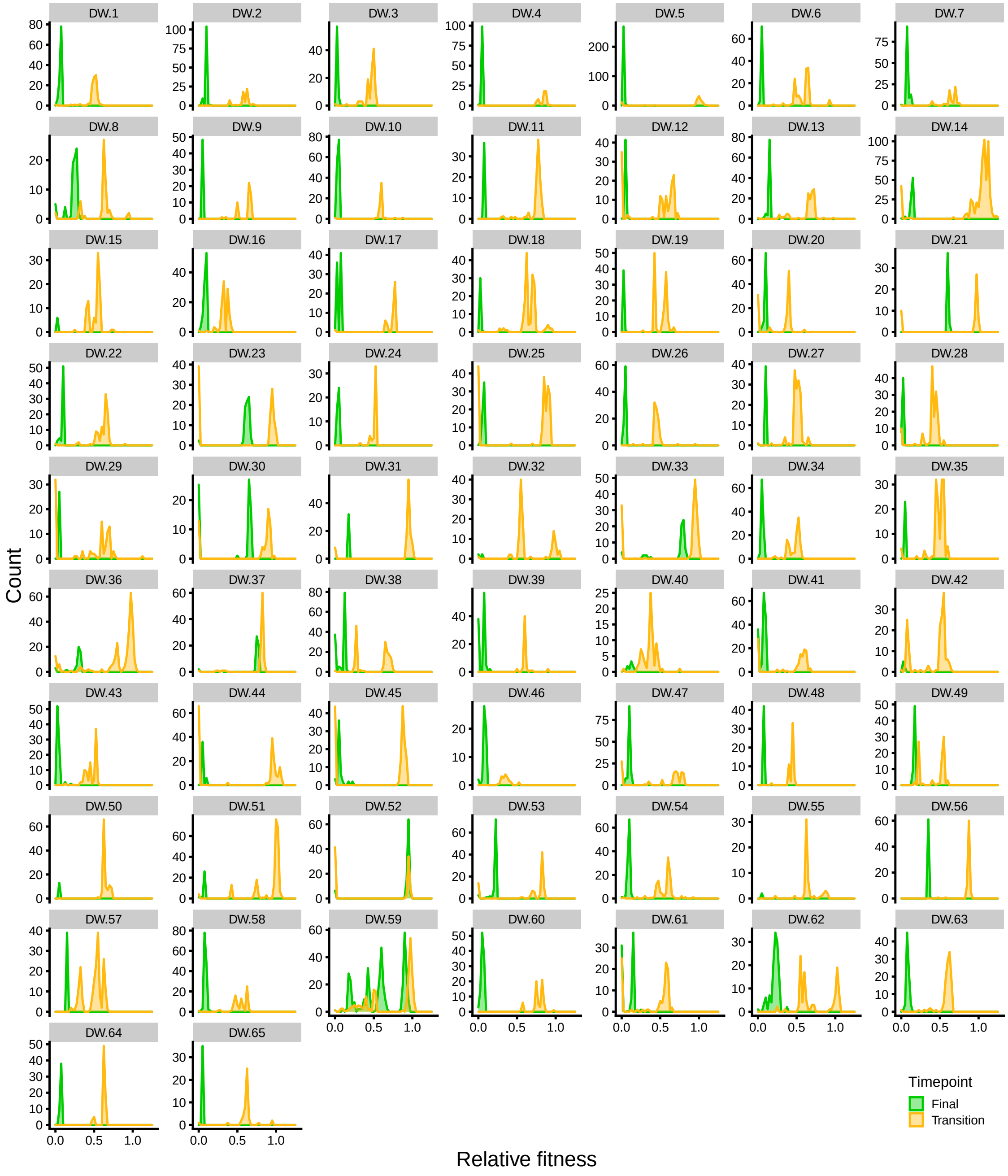


Figure S6: **Distribution of fitness effects of unicellular reversion mutations from the "Dirty Work" experiment at the transition and final time points.** The empirical distribution of fitness effects of reversion mutations for each lineage that evolved multicellularity in our "Dirty work" experiment. Yellow indicates the distribution at the transition point, green at the final time point. Counts are proportional to the adjusted number of unicellular revertants.

#### Fidelity

We directly measure the per site mutation rate ($\mu$) to assess the fidelity of replication in both multicellular and unicellular genotypes. To do this, we record parent and offspring propagule genomes for each replication event that occurs during our growth assays. Each pair of parent and offspring genomes is then aligned using the aphid R package (*10*) and the instruction equivalent of single nucleotide polymorphisms (SNPs) are called wherever there is a mismatch. To restrict our analyses to mutations that could have been the result of either dirty work or the basal mutation rate we record only these SNPs (insertions and deletions are ignored, for example, as they are the result of errors in the copy loop and not from the effects of dirty work). Finally, duplicate mutation calls are removed by comparing SNPs found in offspring derived from the same parent. We repeat this process 100 times for each genotype. We calculate $\mu$ as the total number of unique mutations divided by the number of aligned positions from all offspring genomes. Fidelity is reported as one minus the per site mutation rate (i.e., $1 - \mu$).

#### Effect of time spent evolving as a multicell

In the main text, we treated all lineages that had evolved multicellularity by the end of their evolutionary run equally. However, we observed considerable variation in the time at which multicellularity evolved and this could lead to differences in entrenchment because genomes would spend different periods of time evolving in a multicellular context (*11*). To test this prediction, we examined how several key parameters in our study (entrenchment, number of unicellular revertants, revertant relative fitness, revertant growth rate, and multicell growth rate) changed as a function of the number of generations a population spent evolving as a multicell. Consistent with predictions, entrenchment increased significantly with greater time spent evolving as a multicell (Fig. S7A; Spearman's rank-order correlation, $\rho = 0.3$, $p = 0.02$). Only the change in the number of unicellular revertants did not show a significant correlation with time spent evolving as a multicell (Fig. S7B; Spearman's rank-order correlation, $\rho = 0.08$, $p = 0.53$). The change in unicell revertant relative fitness declined strongly as a function of time spent evolving as a multicell (Fig. S7C; Spearman's rank-order correlation, $\rho = -0.34$, $p = 5.5e-3$) despite the positive correlation betweeen revertant growth rate and time as multicell (Fig. S7D; Spearman's rank-order correlation, $\rho = 0.34$, $p = 6.1e-3$) because the growth rate improvements were vastly outstripped by those of their multicellular progenitor strains (Fig. S7E; Spearman's rank-order correlation, $\rho = 0.86$, $p < 2.2e-16$).

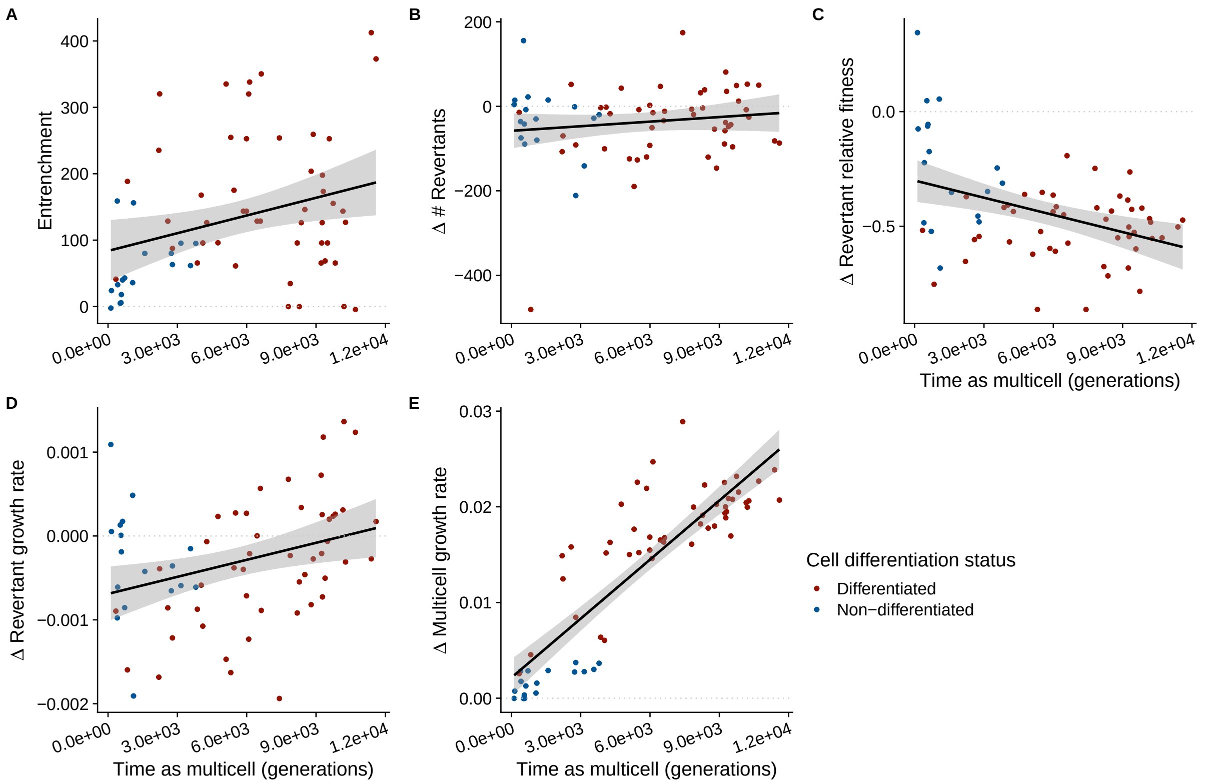


Figure S7: **Effect of time spent evolving as a multicell.** We examine the correlation between time spent evolving as a multicell and the change in five key parameters in our experiment: entrenchment (**A**), number of unicellular revertants (**B**), revertant relative fitness (**C**), revertant growth rate (**D**), and multicell growth rate (**E**). Lineages that evolved germ-soma differentiation appear in red, undifferentiated lineages in blue.

### Distributed Dirt Model

The distributed dirt model disrupts the forces underpinning division of labor by separating the performance of a mutagenic task from its mutagenic consequences. In particular, if a cell within an organism performs a mutagenic task, the performing cell accrues the resource from the task, but the cell with the least amount of mutagenic damage is probabilistically mutated and thus is assigned "the dirt."

The other parameters within the experiment remain exactly the same as the central division of labor experiment. Cells are still able to communicate, perform tasks, tissue accrete, produce propagules, and become propagule-ineligible. Accordingly, the following downstream analyses were conducted in exactly the same way as described for the Dirty Work experiment: calculating entrenchment (Fig. S8), filtering revertants (Figs. S9, S10), and plotting the distribution of fitness effects of reversion mutations (Fig. S11).

Using the combined datasets from the "Dirty work" treatment and the "Distributed dirt" treatment, we also examined the overall correlations between key parameters in our system that affect either entrenchment or unicellular revertant fitness (Figs. S12, S13).

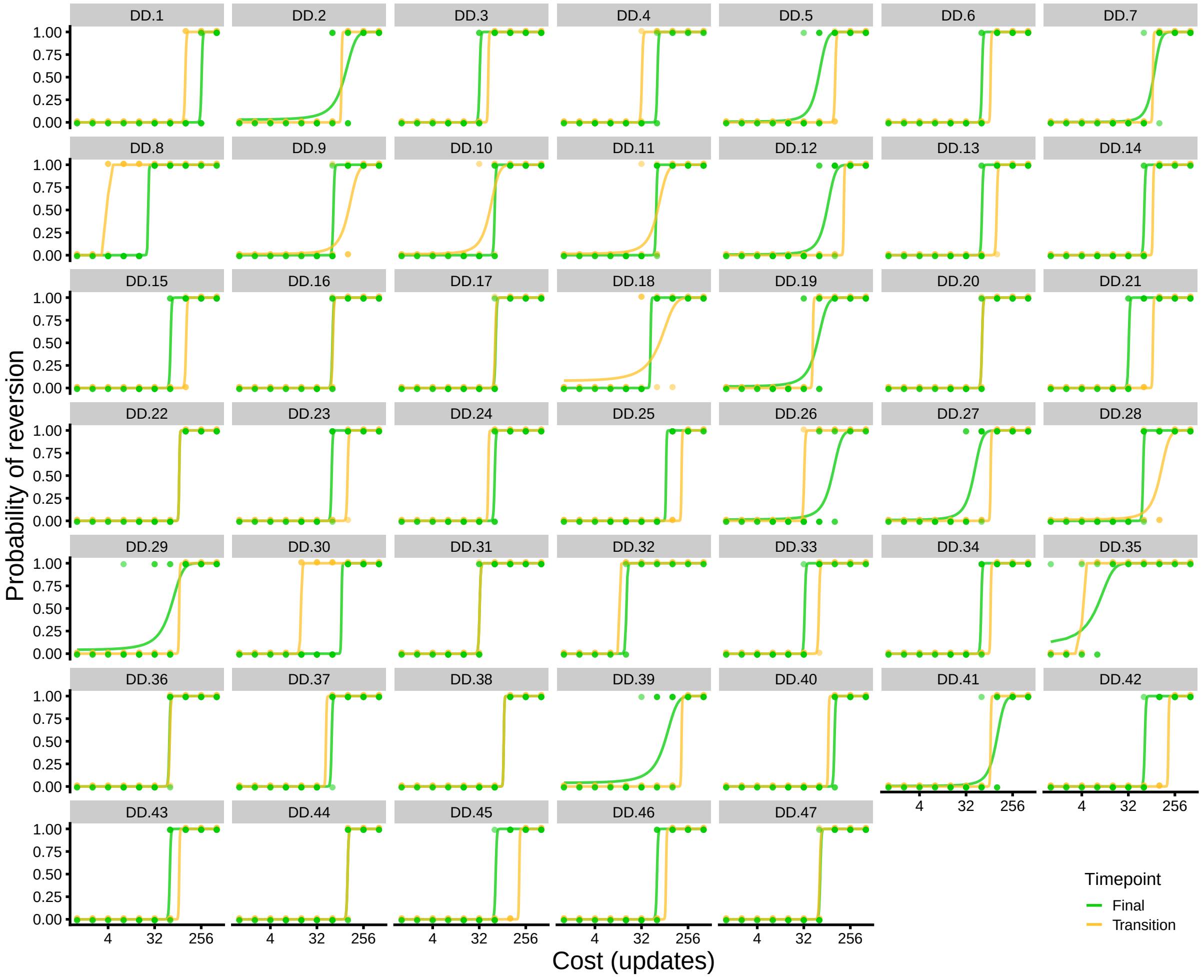


Figure S8: **A comparison of the stability of multicellularity at the transition and final time points of our Distributed Dirt evolution experiment.** The stability of multicellularity was measured for each of the 47 genotypes that evolved multicellularity at the transition (yellow) and final (green) time points of our experiment. The full range of cost values is shown with the outcomes of each replicate plotted as an indicator variable of the dominance of unicellular revertants. Logistic fits of the indicator variable are shown in yellow and green, representing the transition and final time points, respectively.

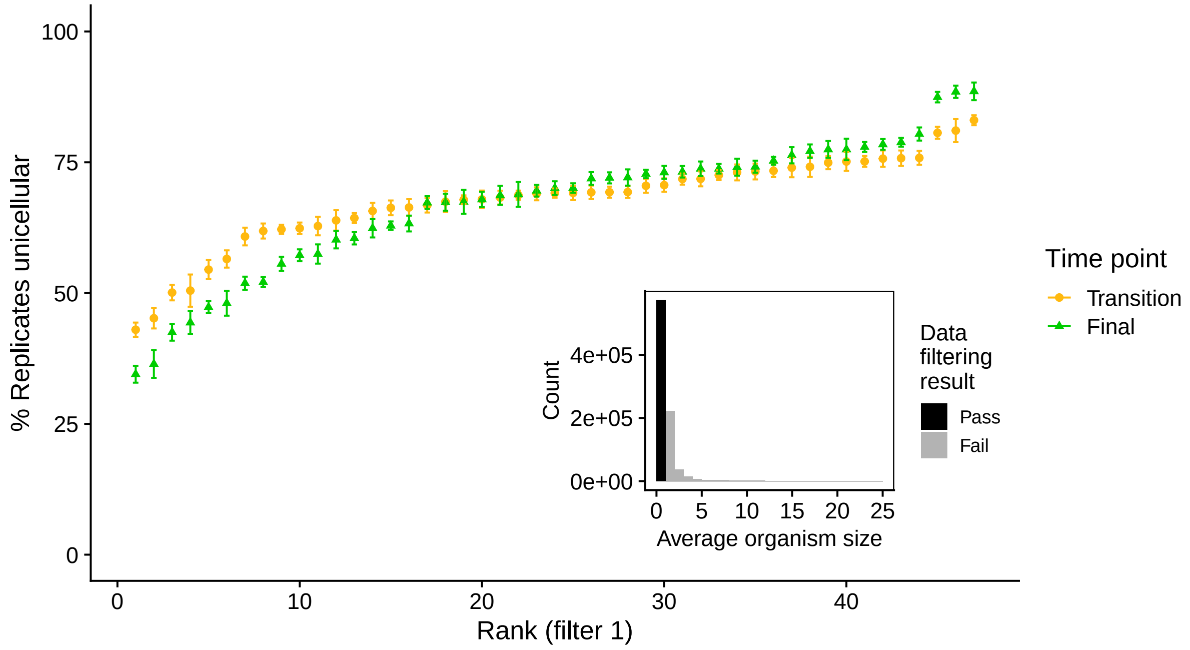


Figure S9: **Filtering unicellular revertants from the "Distributed Dirt" experiment that re-evolve multicellularity.** The mean and standard error for the percent of replicate populations that remain unicellular for the transition (yellow) and final (green) time points of our experiment. Unicellular revertant genotypes are grouped according to their multicellular parental genotypes and ranked from least to most replicates that remain unicelllual for each time point independently. (**Inset**) Schematic representation of our filtering criterion using data aggregated across all genotypes and both time points. We filter out any population in which the average organism size exceeds 1, indicating that multicellularity has re-evolved. Grey color indicates replicates that fail to pass the filter, black indicates those that do.

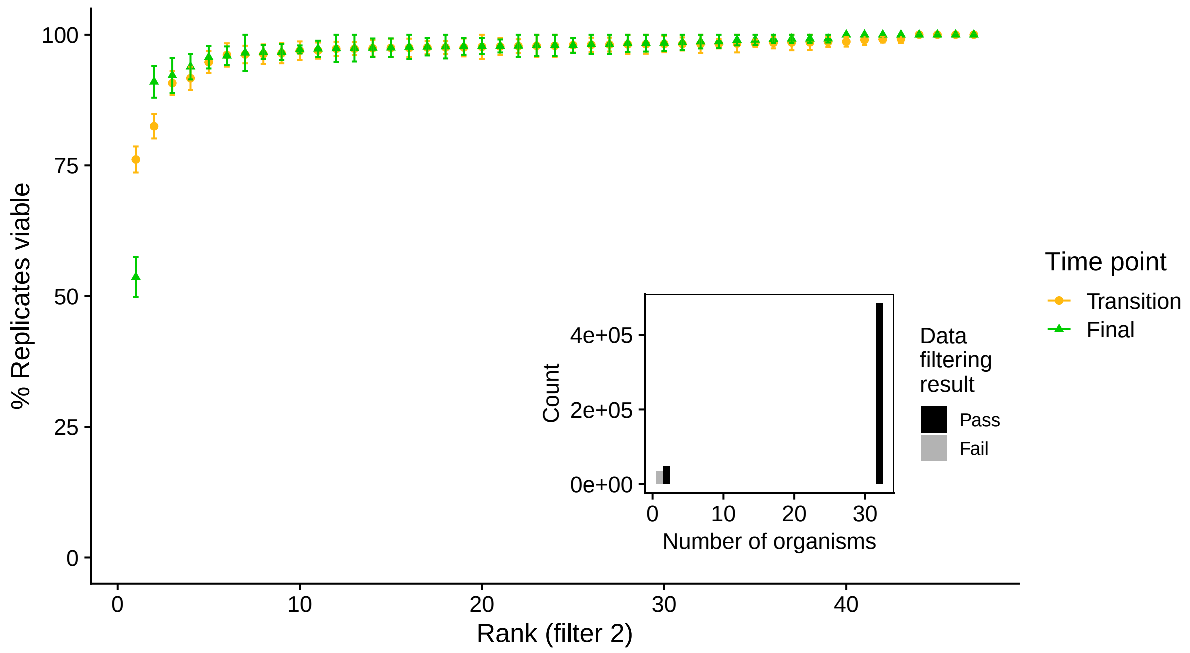


Figure S10: **Filtering unicellular revertants from the "Distributed Dirt" experiment that fail to reproduce.** The mean and standard error for the percent of replicate populations that successfully reproduce for the transition (yellow) and final (green) time points of our experiment. Unicellular revertant genotypes are grouped according to their multicellular parental genotypes and ranked from least to most replicates that remain unicellluar for each time point independently. (**Inset**) Schematic representation of our filtering criterion using data aggregated across all genotypes and both time points. We filter out any population in which the number of organisms at the end of the growth assay is less than 2, indicating that the organism that seeded the population has failed to reproduce. Grey color indicates replicates that fail to pass the filter, black indicates those that do.

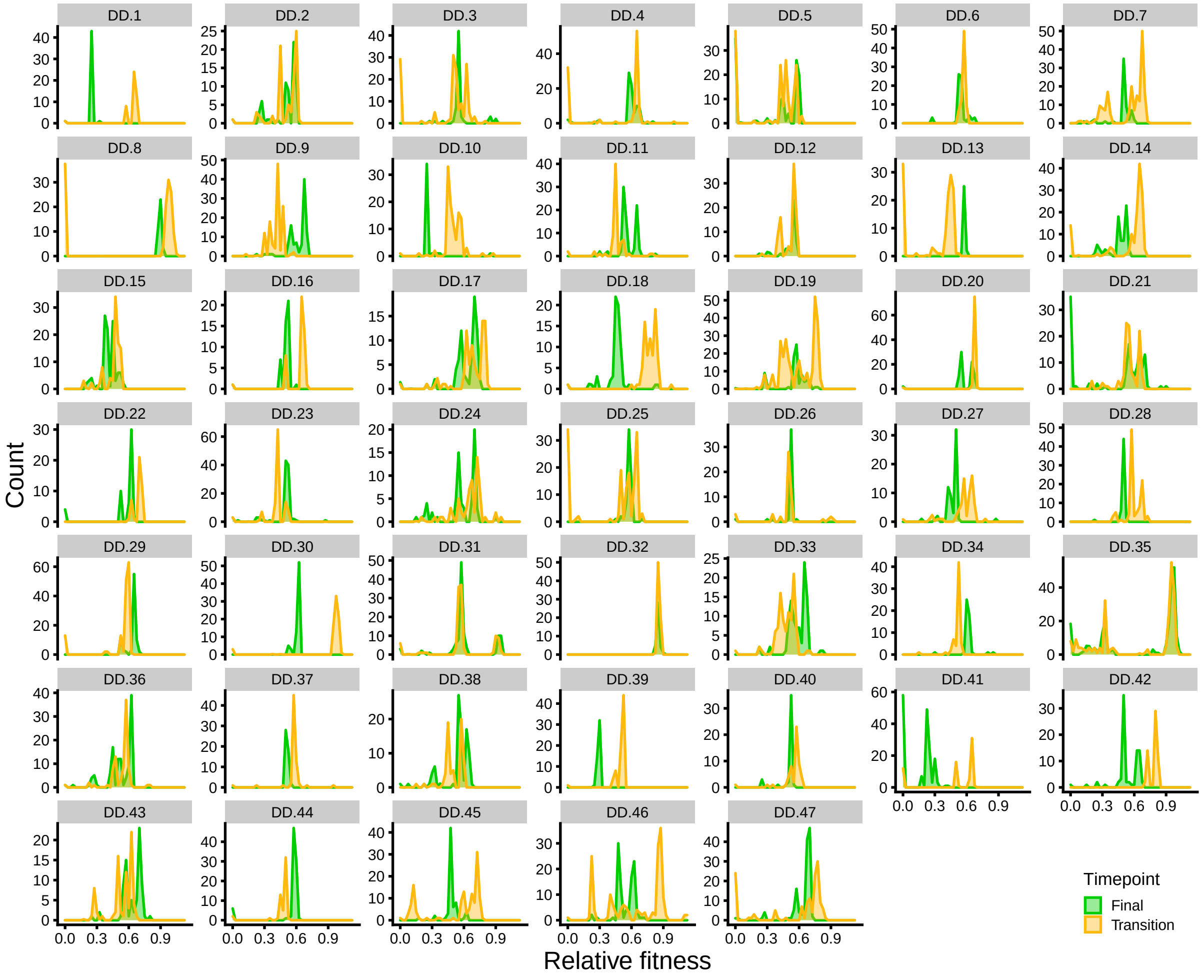


Figure S11: **Distribution of fitness effects of unicellular reversion mutations from the "Distributed Dirt" experiment at the transition and final time points.** The empirical distribution of fitness effects of reversion mutations for each lineage that evolved multicellularity in our "Distributed Dirt" experiment. Yellow indicates the distribution at the transition point, green at the final time point. Counts are proportional to the adjusted number of unicellular revertants.

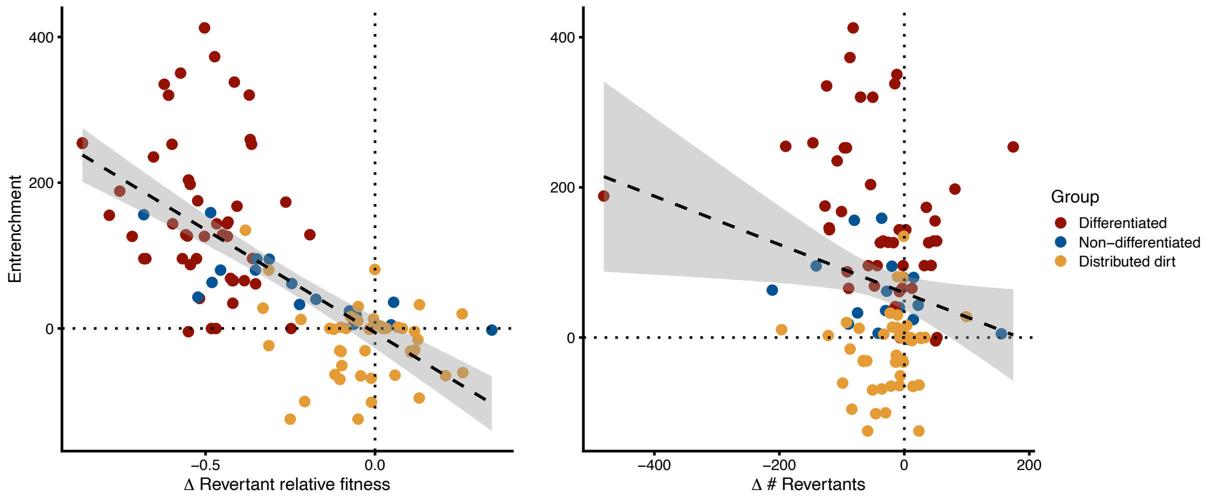


Figure S12: **Entrenchment vs. relative fitness and number of revertants.** We examine the correlation between entrenchment and its two component parameters: revertant relative fitness (**A**) and the number of unicellular revertants (**B**). Differentiated lineages from the dirty-work experiment (red), undifferentiated dirty-work lineages (blue), and the distributed-dirt lineages (gold).

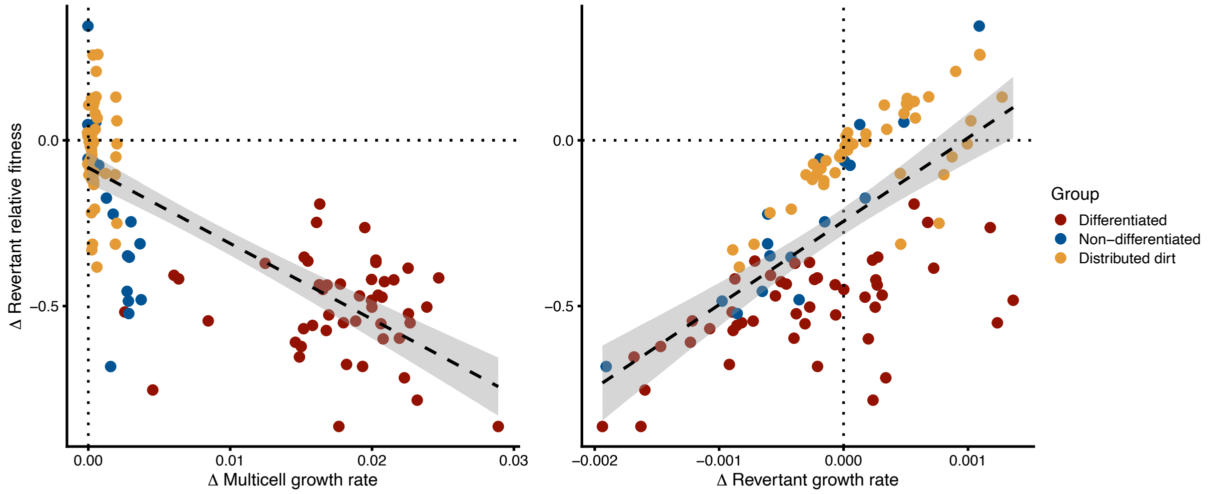


Figure S13: **Relative fitness vs. unicellular growth rate and multicellular growth rate.** We examine the correlation between revertant relative fitness and its two component parameters: the growth rate of the multicellular progenitor strain (**A**) and the growth rate of the unicellular revertant (**B**). Differentiated lineages from the dirty-work experiment (red), undifferentiated dirty-work lineages (blue), and the distributed-dirt lineages (gold).
